# Identification of erythroid cell positive blood transcriptome phenotypes associated with severe respiratory syncytial virus infection

**DOI:** 10.1101/527812

**Authors:** Darawan Rinchai, Matthew B Altman, Oceane Konza, Signe Hässler, Federica Martina, Mohammed Toufiq, Mathieu Garand, Basirudeen Kabeer, Karolina Palucka, Asuncion Mejias, Octavio Ramilo, Davide Bedognetti, Encarnita Mariotti-Ferrandiz, David Klatzmann, Damien Chaussabel

**Affiliations:** Sidra Medicine, Doha, Qatar; Benaroya Research Institute, Seattle, WA, USA; University of Washington, Seattle, WA, USA; AP-HP, Hôpital Pitié-Salpêtrière, Biotherapy (CIC-BTi) and Inflammation-Immunopathology-Biotherapy Department (i2B), Paris, France; Sorbonne Université, INSERM, Immunology-Immunopathology-Immunotherapy (i3), Paris, France; Jackson Laboratory for Genomic Medicine, Farmington, CT, United States; Division of Infectious Diseases, Nationwide Children’s Hospital, Columbus, OH, United States

## Abstract

Biomarkers to assess the severity of acute respiratory syncytial virus (RSV) infection are needed. We conducted a meta-analysis of 490 unique profiles from six public RSV blood transcriptome datasets. A repertoire of 382 well-characterized transcriptional modules was used to define dominant host responses to RSV infection. The consolidated RSV cohort was stratified according to four traits: “interferon response” (IFN), “neutrophil-driven inflammation” (Infl), “cell cycle” (CC), and “erythrocytes” (Ery). Eight prevalent blood transcriptome phenotypes were thus identified. Among those three Ery+ phenotypes comprised higher proportions of patients requiring intensive care. We posit that the erythrocyte module is linked to an overabundance of immunosuppressive erythroid cells that might underlie progression to severe RSV infection. These findings outline potential priority areas for biomarker development and investigations into the immune biology of RSV infection. The approach that was employed here will also permit to delineate prevalent blood transcriptome phenotypes in other settings.

## INTRODUCTION

Respiratory syncytial virus (RSV) infection is the leading cause of hospitalization and the second cause of infant mortality worldwide (1). There are well-characterized populations at risk for severe disease, but most infants who develop a severe RSV infection have no underlying health conditions (2,3). The mechanisms underlying RSV morbidity are poorly understood, but studies suggest that immature or under-developed lungs and/or a dysregulated immune response might have a role (4).

Several groups of researchers, including us, have undertaken blood transcriptome profiling studies of patients with RSV infection (5–13). This approach involves measuring the abundance of blood leukocyte transcripts on a genome-wide scale (14,15). Whole blood comprises a heterogeneous mix of leukocyte populations; thus, changes in transcript abundance might be attributable to either gene expression regulation or relative changes in cell abundance. Regardless, blood transcriptome profiling remains one of the most straightforward approaches to implement in clinical settings and on a large scale (15). Among the RSV blood transcriptome studies, several aimed to identify factors associated with severe disease. For example, we reported an increase in abundance of neutrophil, inflammation and erythrocyte genes in severe pediatric cases (7). Brand et al. pinpointed that an increase in abundance of transcripts coding for Olfactomedin 4, a factor involved in inflammatory responses, is strongly associated with disease severity (10). More recently, Do et al. linked RSV disease severity with follicular T helper cell development and BCL6-dependant inflammation (9).

Building consensus around biomarker signatures and finding a path to clinical utility may involve performing meta-analyses that regroup datasets derived from multiple independent studies (16). Work that permitted the development of novel diagnostic products for sepsis provides a good example (17,18). An obvious benefit of consolidating data from multiple studies is that it permits to achieve larger sample sizes. Arguably the heterogeneity of the patient populations and clinical settings could also help improve the robustness of the resulting biomarker signatures (16). Challenges include the presence of important technical variability between studies, such as in the sampling methods, the profiling platform used or data pre-processing. Another potential limitation is the varying depth and lack of harmonization of sample or subject information available from one study to another.

Such meta-analyses tend to focus on the identification of consensus biomarker signatures: here, the deliverable is a set of differentially expressed genes or predictors of a given clinical outcome. In the present work we endeavored to identify discrete molecular traits (e.g. “interferon = IFN”, “inflammation = Inf”, “Erythrocytes” = Ery”) underlying inter-individual differences among patients with an RSV infection. We used such traits to define blood transcriptome phenotypes and to stratify patient cohorts on the basis of their individual status of each trait: increased, decreased or unchanged vs the uninfected comparators (e.g. of a given phenotype being IFN+ Infl0 Ery-). The next key step was to assess the clinical relevance of such a classification, for instance in terms of differences in the degrees of RSV disease severity. We finally endeavored to investigate the biological basis of the inter-individual variation being measured, in particular for the traits showing the highest degree of association with severe presentations.

A fixed repertoire of transcriptional modules formed the basis for this work. This repertoire consists of a collection of co-expressed gene sets. Co-expression was determined in a collection of reference datasets encompassing 16 distinct immunological states (19) (see methods section). This 382-module repertoire is “fixed”, in the sense that it serves as a reusable framework for analysis and interpretation of transcriptome data. As such, transcriptional modules are not re-formed every time a new dataset is analyzed. Using transcriptional modules is a key aspect of our approach. Indeed, it is the drastic reduction in the number of variables that permits the selection of molecular traits that underpin patient phenotyping and cohort stratification. The fact that this repertoire is fixed is also important, as it permits considerable functional annotations of these transcriptional modules. This annotation can in turn prove critical in unravelling the biological significance of the patient phenotypes being defined.

In summary, we present here a meta-analysis of six RSV blood transcriptome datasets that include 490 unique subject profiles. Specifically, we aimed to: 1) measure inter-individual variability and molecularly stratify RSV patients; 2) identify the associations between patient molecular phenotypes, clinical parameters and outcomes; and 3) identify and interpret the immunobiological processes associated with each molecular phenotype.

## RESULTS

### A collection of RSV blood transcriptome datasets can be assembled from earlier submissions to public repositories

Several researchers investigating the host responses to RSV infection have made their blood transcriptome datasets public. We consolidated the datasets contributed by six independent studies (5–7,20–22) and performed meta-analyses to delineate distinct blood transcriptome phenotypes among RSV subjects. A criterion for including studies in this meta-analysis was the availability of uninfected controls. This point is important because control groups serve as a common denominator between studies and provide the basis for data normalization. Thus, public datasets for which such controls were unavailable could not be included in this meta-analysis.

As each study had different goals and designs, it was first important to identify the key differences, so that they are accounted for when interpreting the meta-analysis results. Information about the six studies is summarized in **Table 1**. The most notable outlier in this collection was the dataset from Liu et al. [GSE73072 (21)] as it consisted of samples from adult subjects collected before and after experimental exposure to RSV. All other studies comprised pediatric subjects with community-acquired RSV infection and a separate group of uninfected controls. Among the latter, the study by Mejias et al. addressed the question of disease severity most directly [GSE38900 (7)], while the work of de Steenhuijsen Piters [GSE77087 (6)] examined the effects of of microbiome composition on the disease course and blood transcriptome signatures. Rodriguez-Fernandez et al. examined the influence of RSV genotypes on blood transcriptional signatures [GSE103842 (5)]]. The study from McDonald et al. focused on identifying pathways involved in disease pathogenesis [GSE80179 (22)]. In the study by Herberg et al. [GSE42026 (20)] the RSV dataset was mostly used as a comparator in a study focusing on responses to H1N1 influenza. The latter two studies were conducted in Europe, while all others were conducted in the United States. Finally, in terms of technical variables, samples from the adult exposure study were run using Affymetrix GeneChips, while the others were run on Illumina BeadArrays. The sample types were otherwise homogenous across all studies and consisted of RNA stabilized whole blood. The studies used one of two popular commercial sample collection tubes for this type of application: Paxgene blood RNA tubes (3 studies) or Tempus tubes (3 studies).

**Table 1:** Description of the public RSV blood transcriptome datasets.

| GEO ID | Reference | Title | N Subjects | Age demographics | Sample type | Platform | Cluster # (Figure 1A) |
| --- | --- | --- | --- | --- | --- | --- | --- |
| **GSE42026** | Herberg et al. / 23901082 / (20) | Transcriptomic profiling in childhood H1N1/09 influenza reveals reduced expression of protein synthesis genes | 22 RSV & 33 controls | Pediatric | Whole blood (PaxGene) | Illumina HumanHT-12 V3.0 | 1 |
| **GSE38900** | Mejias et al. / 24265599 / (28) | Genome-wide analysis of whole blood transcriptional response to RSV, Influenza and Rhinovirus (LRTI) in children | 107 RSV & 31 controls | Pediatric | Whole blood (Tempus) | Illumina Human HT-12 V3.0 ] | 1 |
| **GSE80179** | McDonald et al. /27822537 / (22) | A simple screening approach to prioritize genes for functional analysis identifies a role for IRF7 in the control of RSV disease. | 27 RSV & 52 controls | Pediatric | Whole blood (PaxGene) | Illumina HumanHT-12 V4.0 | 1 |
| **GSE73072** | Liu et al. / 26801061 / (21) | Host gene expression signatures of H1N1, | 20 subjects 2 time points | Adult | Whole blood (PaxGene) | Affymetrix GeneChip Human | 2 |
|  |  | H3N2, HRV, RSV virus infection in adults |  |  |  | Genome U133A 2.0 Array |  |
| **GSE77087** | de Steenhuijsen Pijters et al. / 27135599 / (6) | Nasopharyngeal microbiota, host transcriptome and disease severity in children with RSV infection | 81 RSV & 23 controls | Pediatric | Whole blood (Tempus) | Illumina HumanHT-12 V4.0 | 2 |
| **GSE103842** | Rodriguez-Fernandez et al. / 29045741 / (5) | RSV Genotypes and Disease Severity in Young Children Hospitalized with Bronchiolitis | 62 RSV & 12 controls | Pediatric | Whole blood (Tempus) | Illumina HumanHT-12 V4.0 | 2 |

RSV: respiratory syncytial virus, LRTI: lower respiratory tract infection

Altogether the consolidated dataset collection that was constituted for this meta-analysis encompassed 490 profiles, of which 319 were from subjects with an RSV infection. We hypothesized that this expanded sample size should permit us to define blood transcriptome phenotypes and stratify patient cohorts more effectively than each individual study could.

### Comparing the changes in transcript abundance across module aggregates identifies a consensus RSV signature

Extracting meaningful biological information from large-scale datasets is a notable challenge. Meta-analyses are particularly compounded by the high degree of technical variation existing between independent studies. To at least partly address such challenges, we used a fixed repertoire of transcriptional modules (19) as a framework for data analysis and interpretation. In brief, this repertoire comprises 382 modules, each formed by a set of genes grouped together based on patterns of co-expression across a reference collection of 16 blood transcriptome datasets. This collection was comprised of 985 unique transcriptome profiles and spanned 16 different immunologically relevant pathological or physiological states. A higher degree of organization was further achieved by organizing, in turn, the modules into 38 “aggregates” (designated A1-A38). This grouping was based on similarities in patterns of transcript abundance, determined this time at the module-level and across the 16 reference datasets. Each aggregate comprised between 1 and 42 modules (27 of the 28 aggregates comprised 2 modules or more).

Here, we mapped the changes in transcript abundance for each dataset against this modular framework. From a practical perspective, this means determining for a given module the percentage of its constitutive transcripts that are significantly changed. This procedure is repeated in turn for each module and across each one of the six datasets comprised in our collection. This approach made it possible to assess, as a first step, the degree of consistency in the RSV response signatures across the datasets. To facilitate interpretation, we represented the changes at the least granular level by showing on a heatmap the abundance profiles for each of the six RSV datasets (columns) across 27 module aggregates (rows) (Figure 1A). From this highly reduced set of variables, we could pinpoint the most conserved molecular signatures across the six datasets. These included seven aggregates showing consistent increases in transcript abundance (observed in at least 5/6 datasets: A26, A27, A28, A33, A35, A37, A38) and two aggregates showing consistent decreases (A1, A3). Changes were also observed for another set of modules but in only 3/6 RSV datasets (A15, A16, A29, A30, A34, A36). Technical or biological parameters (Table 1) did not yield an obvious explanation for the differences between these two groups of studies. The amplitude of the changes in other aggregates was minimal. Taken together, this step permitted the mapping of transcriptional changes measured across different RSV data using the same transcriptional module framework. This was useful in relating changes observed between the studies and pinpointing signatures that appear to be most robustly associated with RSV infection.

**Figure 1:**
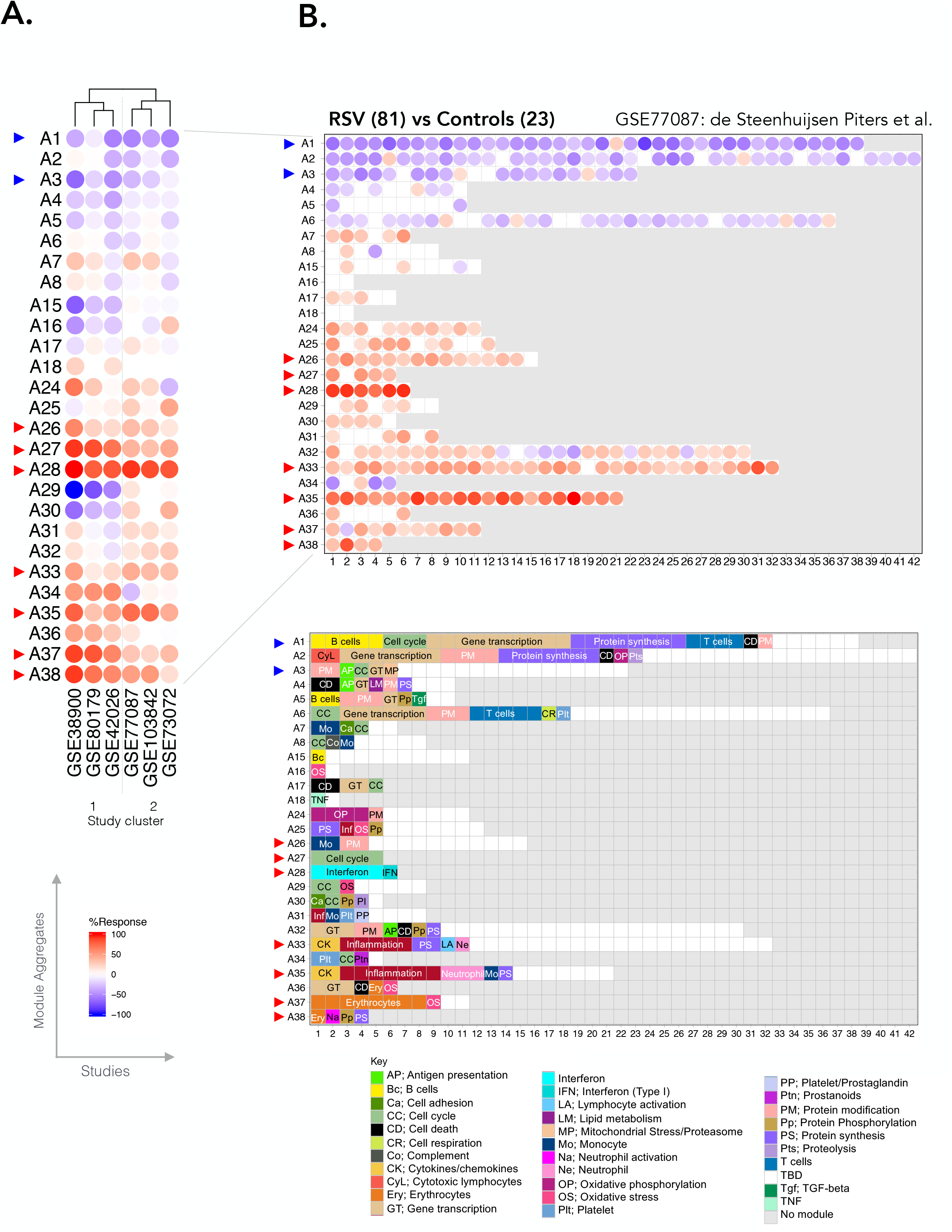
Modular repertoire changes in patients with RSV infection vs uninfected controls. A. Fingerprint heatmap comparing the module aggregate-level patterns of transcript abundance across six RSV datasets. The summarized module aggregate-level values on this heatmap are arranged in rows and the datasets in columns. The datasets are grouped via hierarchical clustering, according to similarities in patterns of transcript abundance across module aggregates. B. Fingerprint grid for GSE77087. Modules are assigned a fixed position on the grid, with each row corresponding to a “module aggregate” constituted of modules following similar patterns of change in transcript abundance. The numbers of constitutive modules for each aggregate range from two (A16) to 42 (A2). Aggregates comprising a single module are not represented on this map (A9-A14; A19-A23). The percentage of constitutive transcripts for a given module showing an increase in abundance in RSV patients over controls is indicated by a red spot. The percentage of constitutive transcripts showing a decrease in abundance for a given module is indicated by a blue spot. The color key at the bottom indicates the functions that have been associated with some of the modules on the grid.

### Changes in transcript abundance can be mapped to a fixed transcriptional module repertoire to facilitate functional interpretation

To functionally interpret the conserved signatures observed across RSV datasets, a more granular level of information is needed. We thus examined transcriptomic changes at the level of the modules forming the 27 aggregates mentioned above. We represented the changes in transcript abundance as grid plots for each of the RSV datasets (**Figure 1B**, **Supplementary Figure 1**). A first vertical reading of the grid across the rows provides a sense of the changes at the aggregate level already summarized in the heatmap that was presented earlier (**Figure 1A**). A second horizontal reading across the columns provides a sense of the changes occurring at the module level within each of the aggregates.

Because the positions on the grid are fixed, it is possible to overlay other information, such as functional annotations (color-coded grid in **Figure 1B**). We found that some of the conserved signatures that were increased during RSV infection comprised modules preferentially associated with interferon responses (A28), inflammation (A33, A35), erythrocytes (A37, A38) and cell cycle (A27), while those that were decreased were associated with lymphocytic responses (A1, A3). Some of these responses are further interpreted below, and all details are accessible via interactive web presentations for modules constituting each aggregate (web links are listed in **Table 2**). The presentations include reports from functional profiling analyses carried out using different tools. Heatmaps representing the patterns of abundance for transcripts constituting each module across reference datasets are also available. Furthermore, a dedicated web application was develop in support of the work presented here and permits users to access the fingerprint grid plots presented here and generate other types of plots which are presented throughout this manuscript. This resource can be accessed via this link: https://drinchai.shinyapps.io/RSV_Meta_Module_analysis/. A video demonstration can be accessed here: https://youtu.be/htNSMreM8es.

**Table 2:**
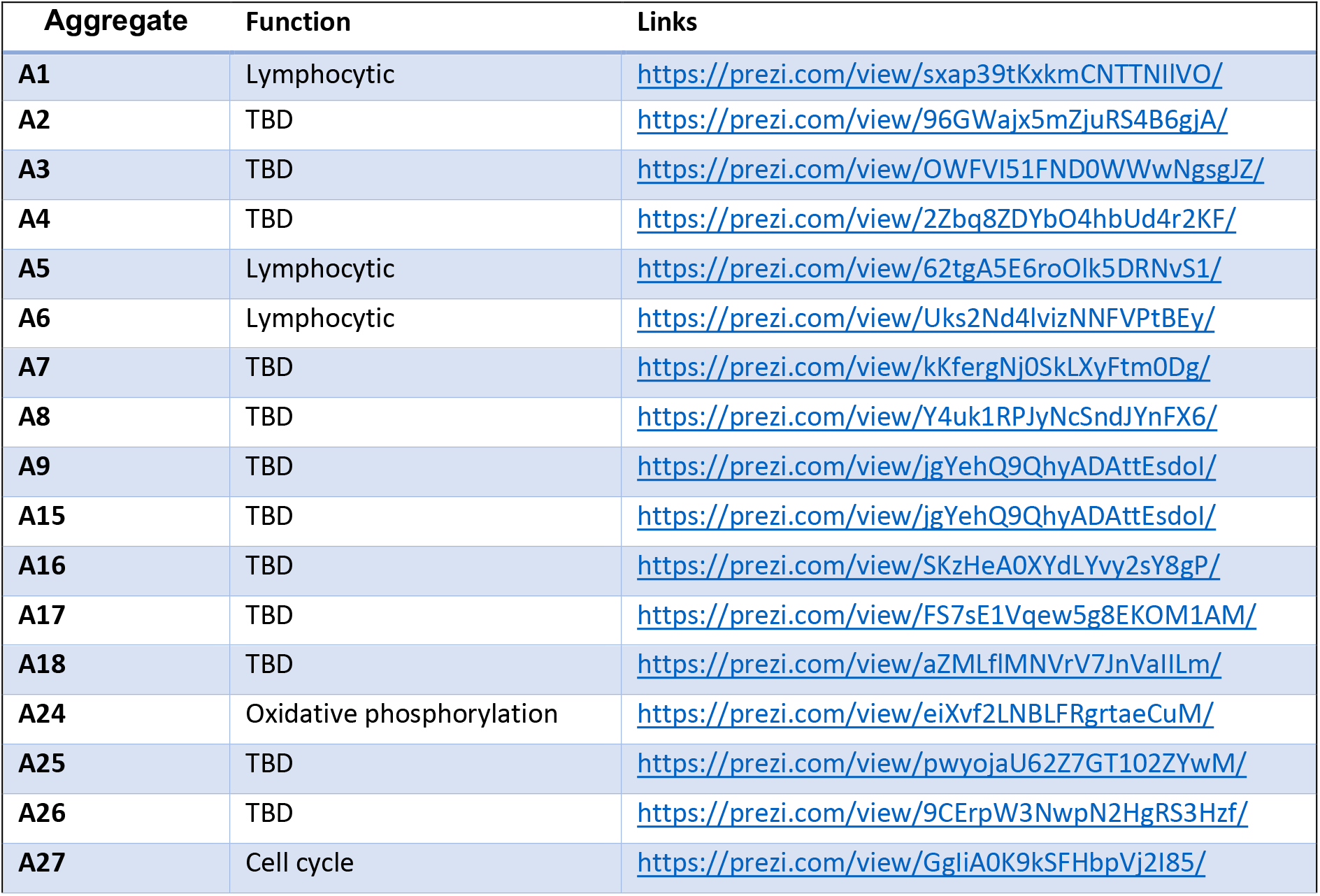
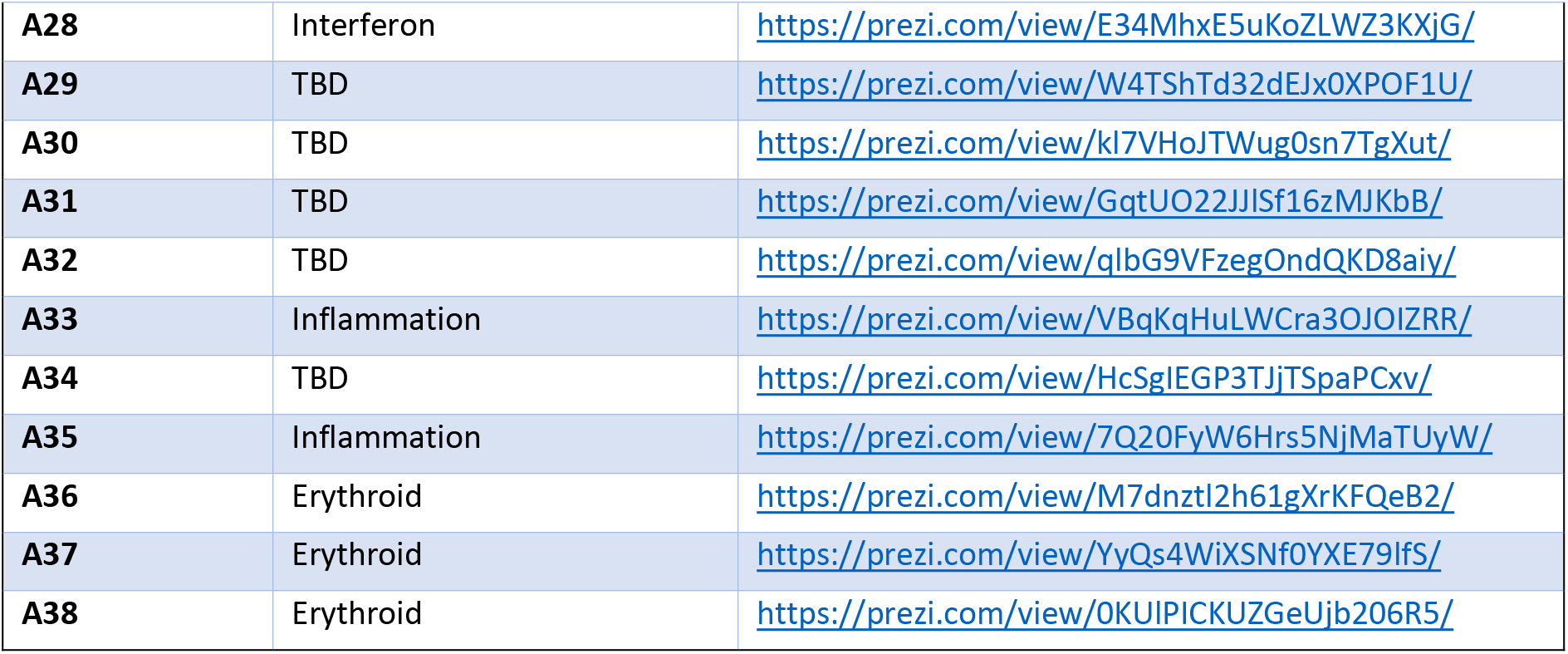
Links to module aggregates annotation pages.

In summary, we mapped the conserved RSV signatures identified to a well-annotated modular framework. This mapping made it relatively straightforward to assign each signature to a predetermined functional category. The added granularity and available online resources make it possible to further dissect those signatures in subsequent analyses, as exemplified below.

### Blood transcriptional signature present a high level of inter-individual among cohorts of RSV patients

Blood transcriptome profiling provides a means to measure inter-individual differences with a high degree of resolution. Understanding the biological and clinical significance of this variability is important but requires the study of large patient cohorts. The collective re-analysis of the six datasets assembled here provides a unique opportunity to investigate inter-individual differences among patients with RSV infection at the molecular level.

The approach that we used next to map changes in transcript abundance for individual patients against the repertoire of transcriptional modules is very similar to that described above for groups of patients. We expressed the changes for each individual RSV patient as a percentage of constitutive genes for which the abundance was increased or decreased compared to the respective control group (see methods for details). As an illustration, we generated a heatmap (**Figure 2**) of the results obtained for subjects comprising the de Steenhuijsen Piters dataset (**Figure 1B**). The patterns in the changes in abundance are only shown for the modules constituting the nine aggregates deemed to be conserved across the collection of RSV datasets (highlighted in **Figure 1**). We generated similar plots for each of the remaining datasets (**Supplementary File 1**). There are also accessible and customizable via our web application (https://drinchai.shinyapps.io/RSV_Meta_Module_analysis/).

**Figure 2.**
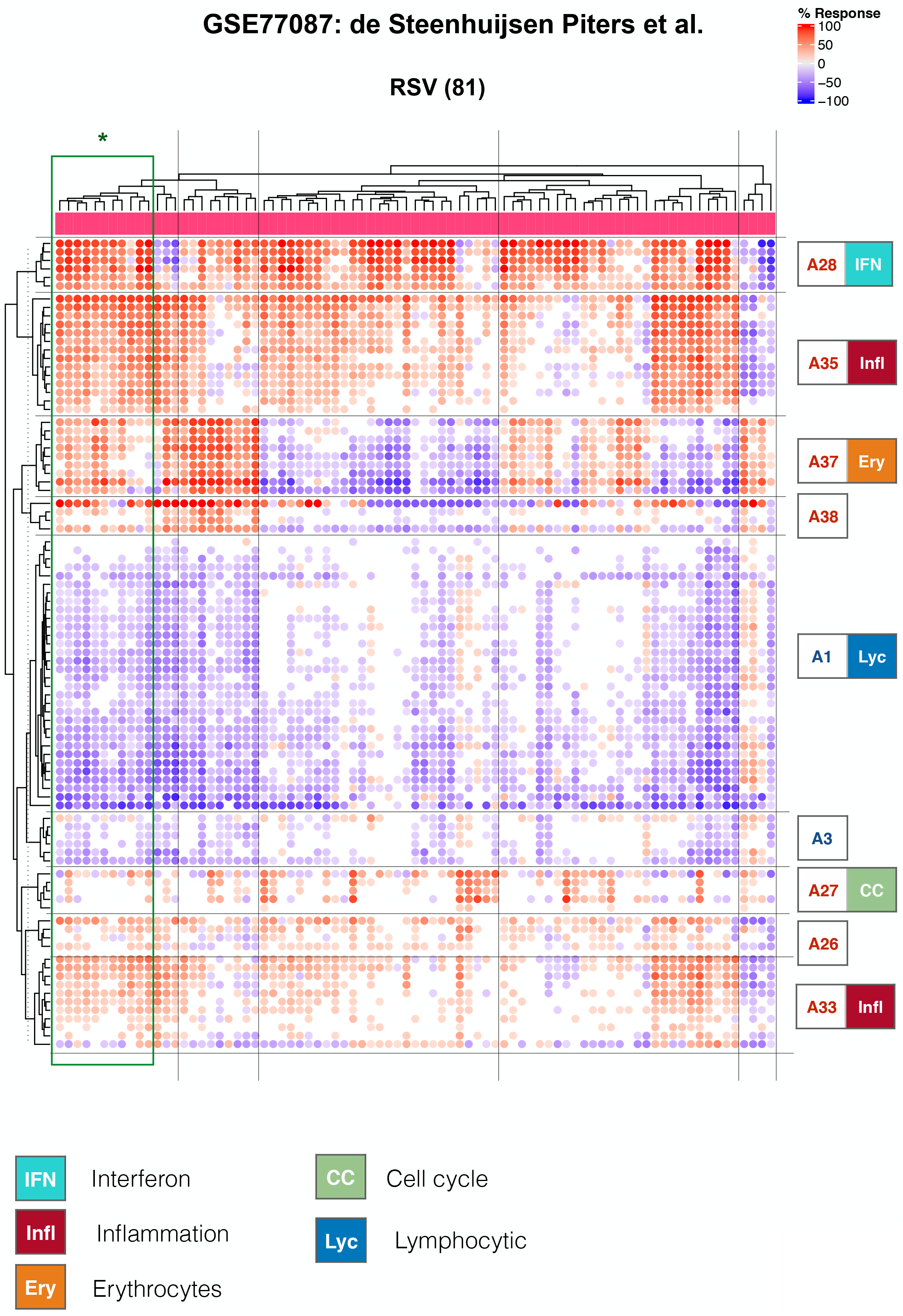
Heatmap representation of changes in abundance of transcriptional modules across RSV infected individuals. This heatmap was generated for the GSE77087 dataset that is also represented as a fingerprint grid plot in Figure 1B. The modules comprised in aggregates identified as being conserved (indicated by the colored triangles in Figure 1A) are arranged as rows and the RSV subjects comprised in this dataset are arranged as columns. The colored spots represent the percentage of transcripts within each module deemed to be differentially expressed (up = red, down = blue). The modules are arranged based on similarities in abundance patterns via hierarchical clustering within each aggregate. A general function is attributed to some of the aggregates, as indicated by the colored symbols and key.

From the heatmaps, we observed that inter-individual variability exists even for signatures that at the group level were well-conserved across the six datasets (**Figure 1A**). In reality, only a minority of patients in this illustrative dataset (11/81) matched the prototypical pattern defined above based on conserved changes observed for nine module aggregates (i.e. A1− A3− A27+ A28+ A33+ A35+ A37+ A38+; **Figure 2**). The degree of inter-individual variability differed from one module aggregate to another. For example, in the modules forming aggregates A1 (Lymphocytic) or A33 (Inflammation), changes in abundance only varied in amplitude without, for the most part, showing an inversion of trends. In other modules, inverted trends were much more common, as exemplified by A37 (Erythrocytes).

Taken together, examining changes in transcript abundance at the level of individuals revealed a significant degree of heterogeneity among cohorts of RSV patients. This paradigm was also true for signatures deemed to be conserved when carrying out comparisons at the group level.

### Distinct blood transcriptome phenotypes are identified among a consolidated cohort of RSV patients

The fact that the consensus disease signature defined earlier was not reflected at the level of individual patients highlighted the need to characterize distinct RSV blood transcriptome phenotypes. For this, we used the combined set of patients from the six public transcriptome datasets. First, we generated PCA plots to evaluate the sources of variance among this composite set of samples (**Figure 3A**). The results indicated the absence of study bias when abundance measures where normalized to the respective control group and reduced to module-level summarized values. This finding was largely confirmed when representing inter-individual differences on a tSNE plot (**Figure 3B**) (23). One dataset did show partial separation from the others, but this only concerned a minority of subjects and could also be attributable to biological sources. The same shift was observed on the PCA plot but only along PC3, which accounts for only 9% of the overall variance.

**Figure 3:**
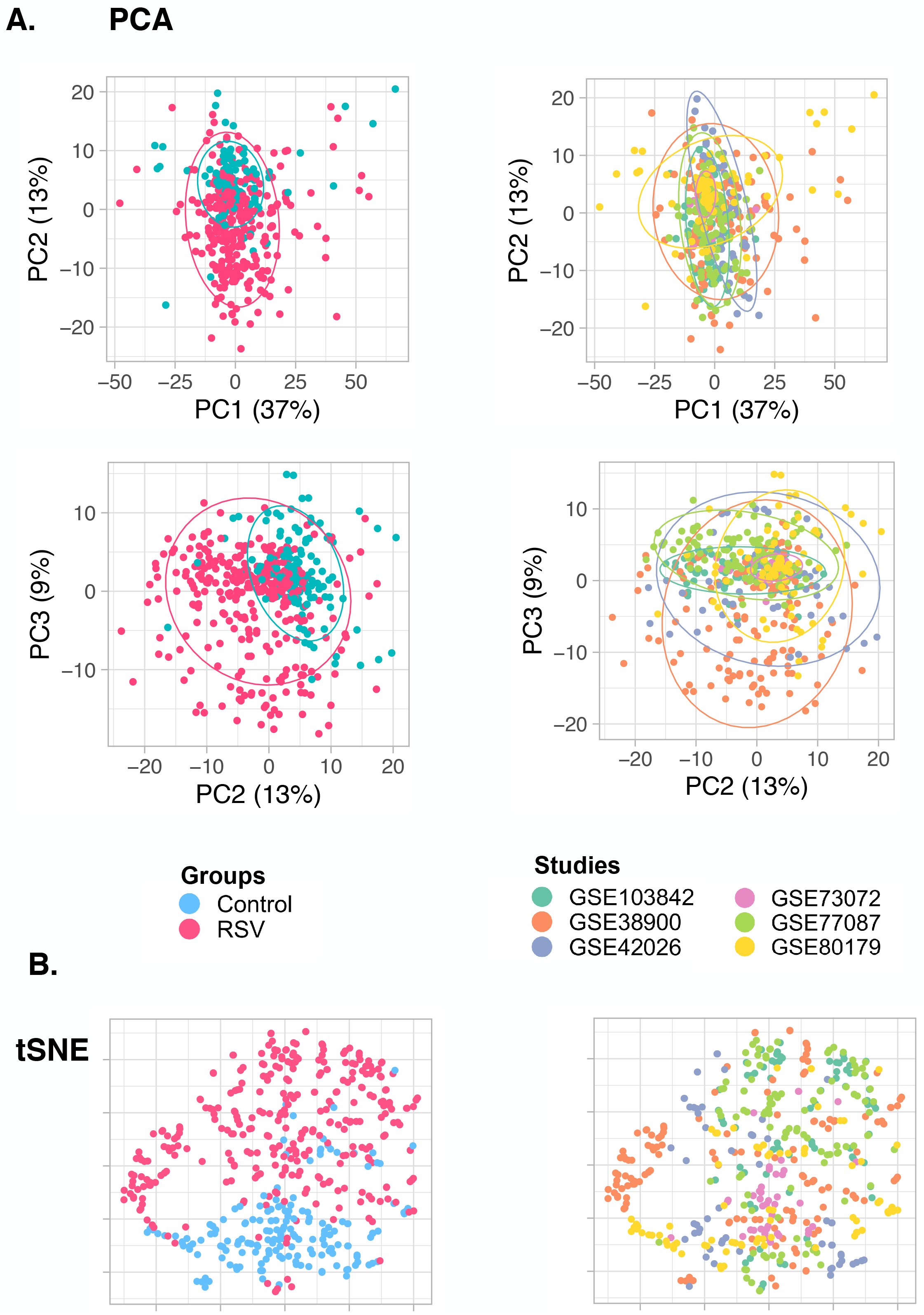
Individual subjects PCA plots and tSNE plots. The modular fingerprint profiles of individual subjects from each of the six different studies were combined and PCA and tSNE plots were generated. The total number of subjects was 490 (319 subjects with RSV infection and 171 controls). **A.** 2D PCA plots. The subjects are color-coded according to grouping information (Groups) and dataset membership (Studies). **B.** tSNE plots. In both types of plot, each sample is represented by a dot. The proximity between the dots is an indication of the modular transcriptional profile similarity.

As a first step, we sought to identify signatures with the highest degree of inter-patient variability, assuming that these would be best able to discriminate patients according to phenotypes. We identified these signatures at the least granular, module-aggregate level, thus starting from a set of only 38 variables (**Supplementary Figure 2**). We selected the following four aggregates: 1) A27, comprising five modules functionally associated with the “cell cycle”; 2) A28, comprising six modules functionally associated with the “interferon response”; 3) A35, comprising 21 modules predominantly associated with “inflammation”; and 4) A37, comprising 11 modules predominantly associated with “erythrocytes”. We discarded other aggregates that exhibited a similar degree of inter-patient variability due to likely redundancy with the four selected aggregates. These aggregates included A33, which like A35 is also associated with “inflammation”, and A38, which like A37 is associated with “erythrocytes”.

Next, we assigned the status for each aggregate signature in a given individual using the corresponding percentage of increased or decreased transcripts: if the value was >15% the aggregate was considered to be increased (noted as +), if ≤15% it was considered to be decreased (noted as −), else it was considered not changed (noted as 0). For example, Infl0/IFN+/CC+/Ery−. We then generated the distribution of subjects constituting the combined RSV cohorts across all 81 possible Infl/IFN/CC/Ery phenotypes (**Figure 4**). We found that a small subset of phenotypes comprised a higher number of patients than others (>10 per phenotype). These phenotypes were all positive for the interferon “trait” (IFN+), positive or showed no changes for the “inflammation” and “cell cycle” traits (Infl+/0 or CC+/0) and exhibited any erythrocytes trait status (Ery+/0/−). Phenotypes where the interferon status was unchanged or decreased were comparatively less prevalent (five patients per group at most), and so were phenotypes where interferon was increased but inflammation or cell cycle phenotypes were decreased.

**Figure 4:**
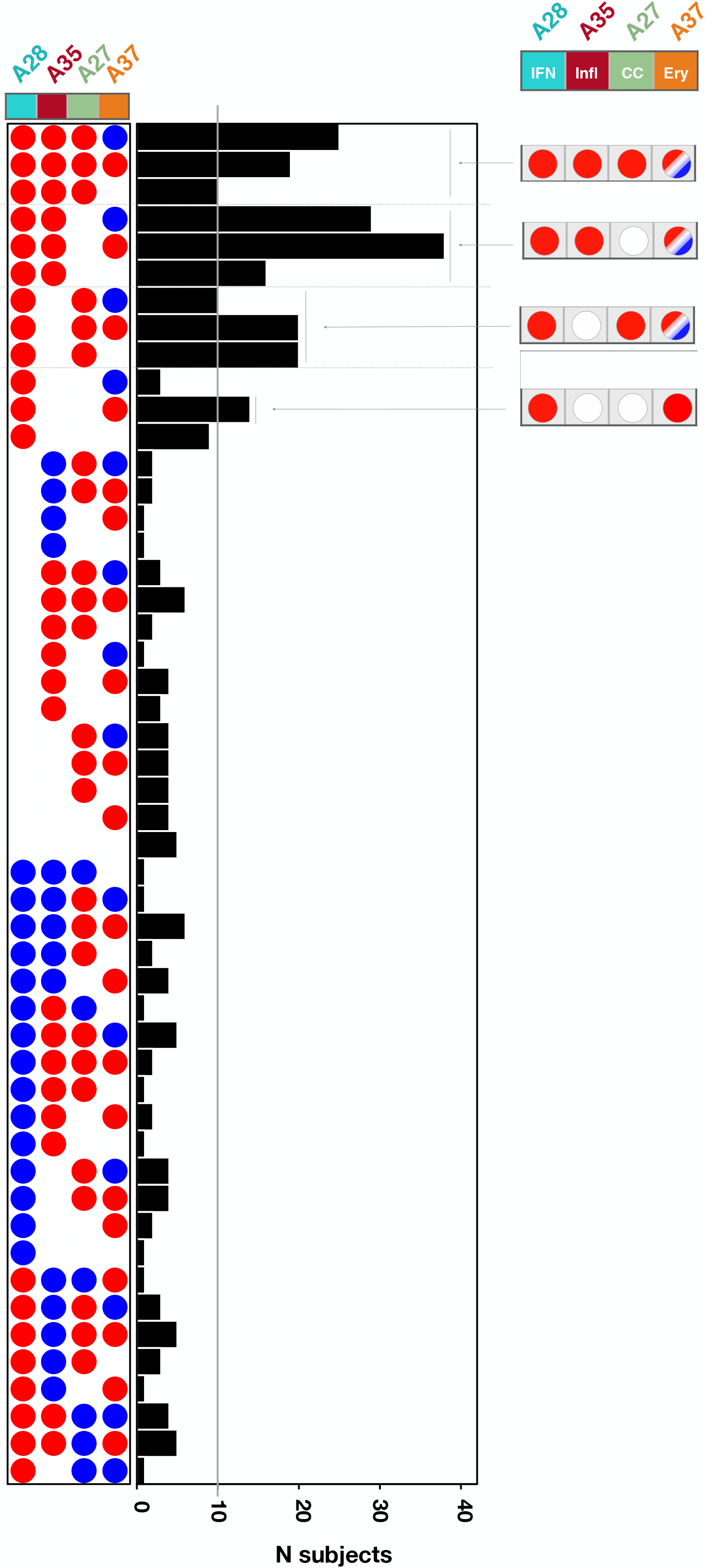
Stratification of RSV patients according to blood transcriptome phenotypes. Phenotypes were defined according to four different “traits”. Each of the 319 RSV patients comprising the consolidated cohort used in this meta-analysis were assigned to a phenotype according to their status for each of the four traits: positive (red), negative (blue) or unchanged (white). This determination was made in reference to each respective control baseline. The bar graph shows the number of patients assigned to each of the phenotypes. The gray line indicates the threshold used to select the phenotypes considered to be the most abundant (< 10 subjects).

Overall, we have shown here that a principled approach using a small subset of highly reduced variables can identify a discrete number of interpretable RSV blood transcriptome phenotypes. It should be noted, however, that alternate classifications could be obtained by modifying the selection criteria. This would be the case if, for instance, more emphasis was put on biological significance than inter-patient variability.

### A subset of RSV blood transcriptome phenotypes is associated with severe disease

An obvious next question is whether the stratification of RSV patients according to blood transcriptome phenotypes, such as those described above, have any clinical relevance. The extent of phenotypic information made available alongside the blood transcriptome datasets varied significantly from study to study. Notably, pertinent information reflecting disease severity (e.g. respiratory rate, transcutaneous O2 saturation) were lacking for many patients. As a result, we had to use a relatively crude metric of disease severity that relied on the type of care the patient required; i.e. whether they were outpatients, inpatients cared for in the ward or were admitted to the pediatric intensive care unit (PICU).

We first visualized the patterns of transcript abundance at the module level for individuals belonging to the seven most prevalent phenotypes (**Figure 5**). This heatmap verified that the phenotypic categories presented a high degree of homogeneity. Upon overlaying the phenotypic information on this plot, we gained a first indication of a possible association between age and disease severity. Specifically, it was possible to discern a trend towards a younger age among Ery+ subjects in comparison to Ery- or Ery0 subjects. Importantly, the different studies also seemed to be well represented in each of the phenotypes, indicative of their underpinnings by biological rather than technical factors.

**Figure 5:**
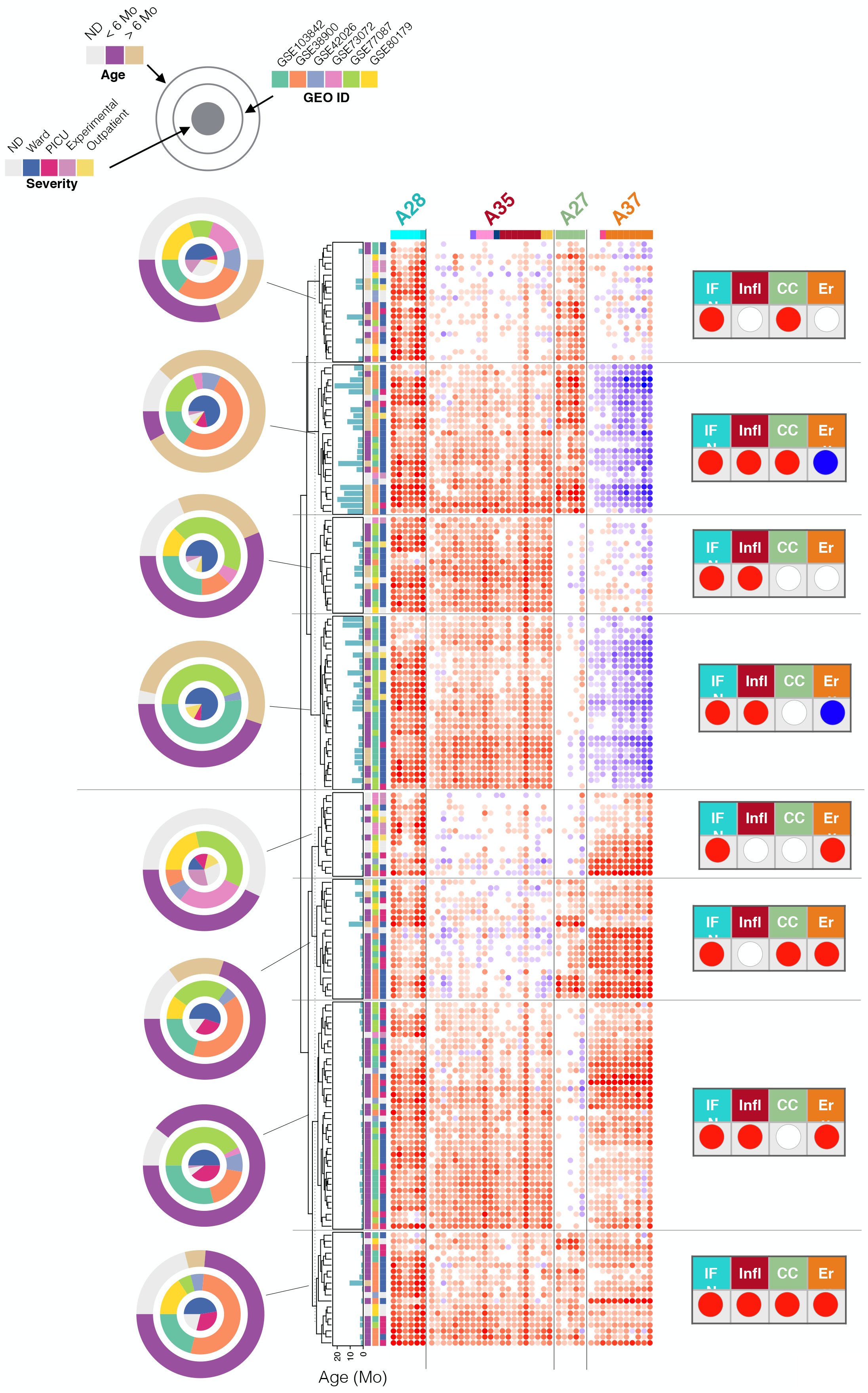
The association of dominant RSV blood transcriptional phenotypes with clinical and demographic attributes. Heatmaps were generated for the seven most prevalent phenotypes identified on the distribution plot in Figure 4. The subjects (rows) were first arranged according to their phenotype, and then arranged within each phenotype according to similarities in abundance patters. Modules constituting the four aggregates selected for the definition of molecular phenotypes are shown as columns. The respective traits for module aggregates A28, A35, A27 and A37 are IFN (interferon), Infl (inflammation), CC (cell cycle) and Ery (Erythrocytes). The status for each phenotype is indicated by a red, blue or white spot (increase, decrease or no change, respectively). The concentric circle plots (right) indicate the distribution of patients constituting each phenotype according to age, study membership and RSV severity status.

We then looked at the relative proportion of severe patients for each high-prevalence phenotype and the contributions by the different studies (**Figure 6A**). Four of the eight phenotypes were Ery+, of which three comprised a proportion of PICU patients that was on average 5.6 times higher than the four Ery- and Ery0 phenotypes. The Ery+ patients included the quadruple positive IFN+/Infl+/CC+/Ery+ phenotype with 32% of PICU patients, while its IFN+/Infl+/CC+/Ery-counterpart had 12% of patients. We found no severe patients in the IFN+/Infl+/CC0/Ery0 group. Furthermore, subjects with Ery+ phenotypes were significantly younger than subjects with Ery-phenotypes (p <0.001, **Figure 6B**).

**Figure 6:**
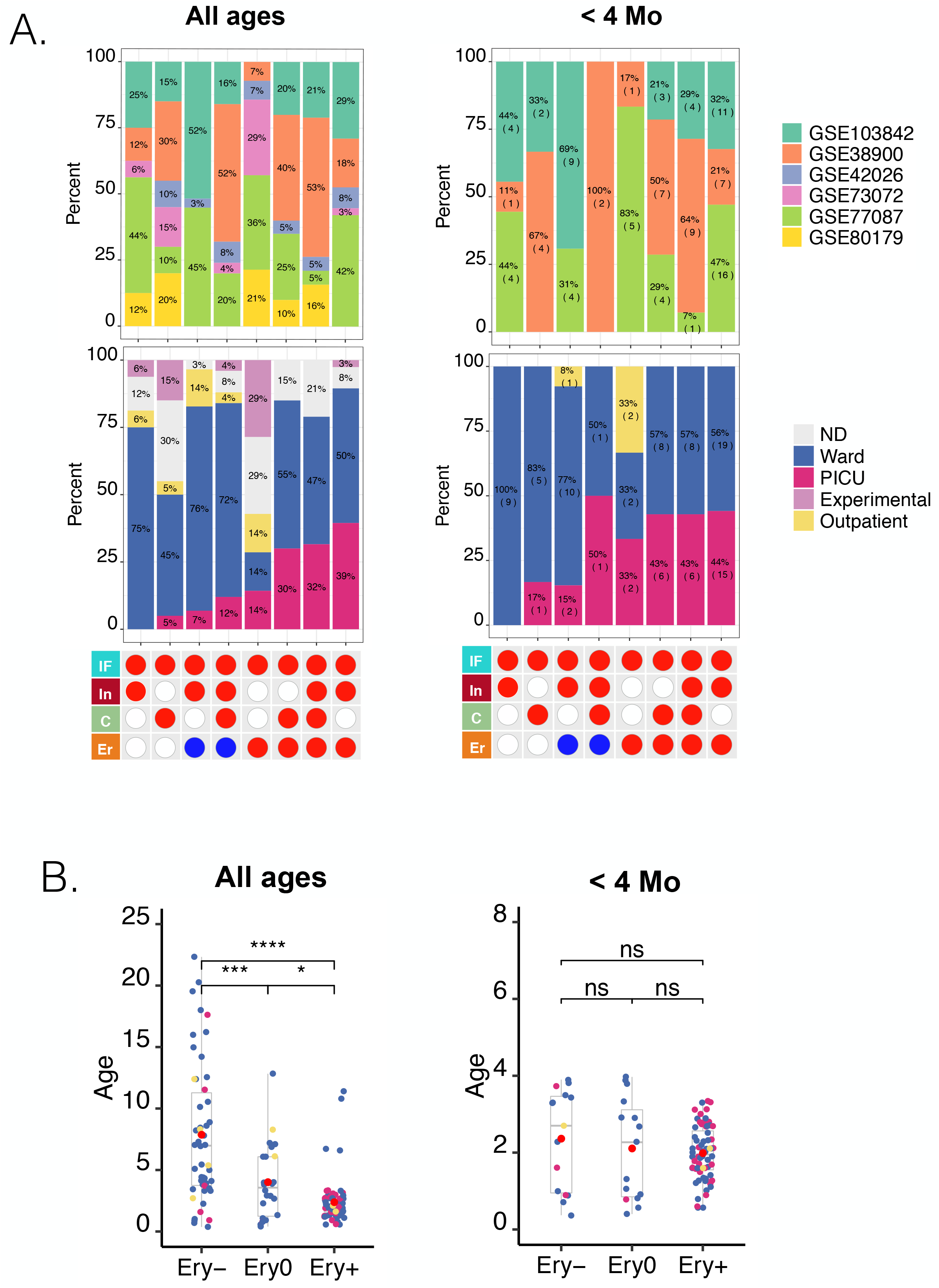
The association of dominant RSV blood transcriptional phenotypes with disease severity as a function of age. A. The relative frequencies of patients in PICU care across dominant IFN/Infl/CC/Ery phenotypes. Bottom panel: the red or blue spots define the status for four traits corresponding to the eight most prevalent RSV phenotypes identified in Figure 4 (<10 subjects). Middle panel: the relative frequency of subjects cared for in the hospital ward, PICU, outpatients, and experimentally exposed subjects. This information was not available for all studies. Top panel: the relative contribution of the different datasets selected for this meta-analysis. The combination of tables and graphs on the left is for all subjects. Information for a subset of subjects <4 months-of-age is shown on the right. B. Ages according to Ery trait status for all patients and a subset aged <4 months old. The box plots represent the age in months of individuals comprising the consolidated RSV cohort used throughout this study. Patients were categorized according to their Ery trait status. The plot on the right shows the same information but of a subset of patients <4 months old. Dots are color-coded according to severity status (Red = PICU, Blue = ward, Yellow = outpatient). * p< 0.05, ** p< 0.01, *** p< 0.001.

We next endeavored to determine whether the presence of the Ery trait was associated with heightened severity, regardless of age. For this we examined the distribution of severe cases across the same phenotypes but focusing on infants <4 months old (**Figure 6A)**. Again, the severe cases were distributed preferentially among Ery+ phenotypes [Ery+ = 29 PICU) cases, Ery−/0 = 4 PICU cases; of note the IFN+/Infl+/CC+/Ery-phenotype comprised only two patients, one of which was a PICU case].

Finally, we investigated the associations between each of the four traits used for RSV patient phenotyping and stratification, and disease severity (**Figure 7**). Here we found that the abundance of transcripts forming the A37/erythrocytes cells aggregate were significantly increased in patients cared in the PICU compared to the ward (Ery trait; p<0.01). We made a similar finding for the A35/“inflammation” aggregate, although to a lesser degree (Infl trait; p<0.05). We found no significant differences were found for the A27/cell cycle or A28/interferon aggregates. Associations can be explored for various aggregates as a function of age via our web application (https://drinchai.shinyapps.io/RSV_Meta_Module_analysis/).

**Figure 7:**
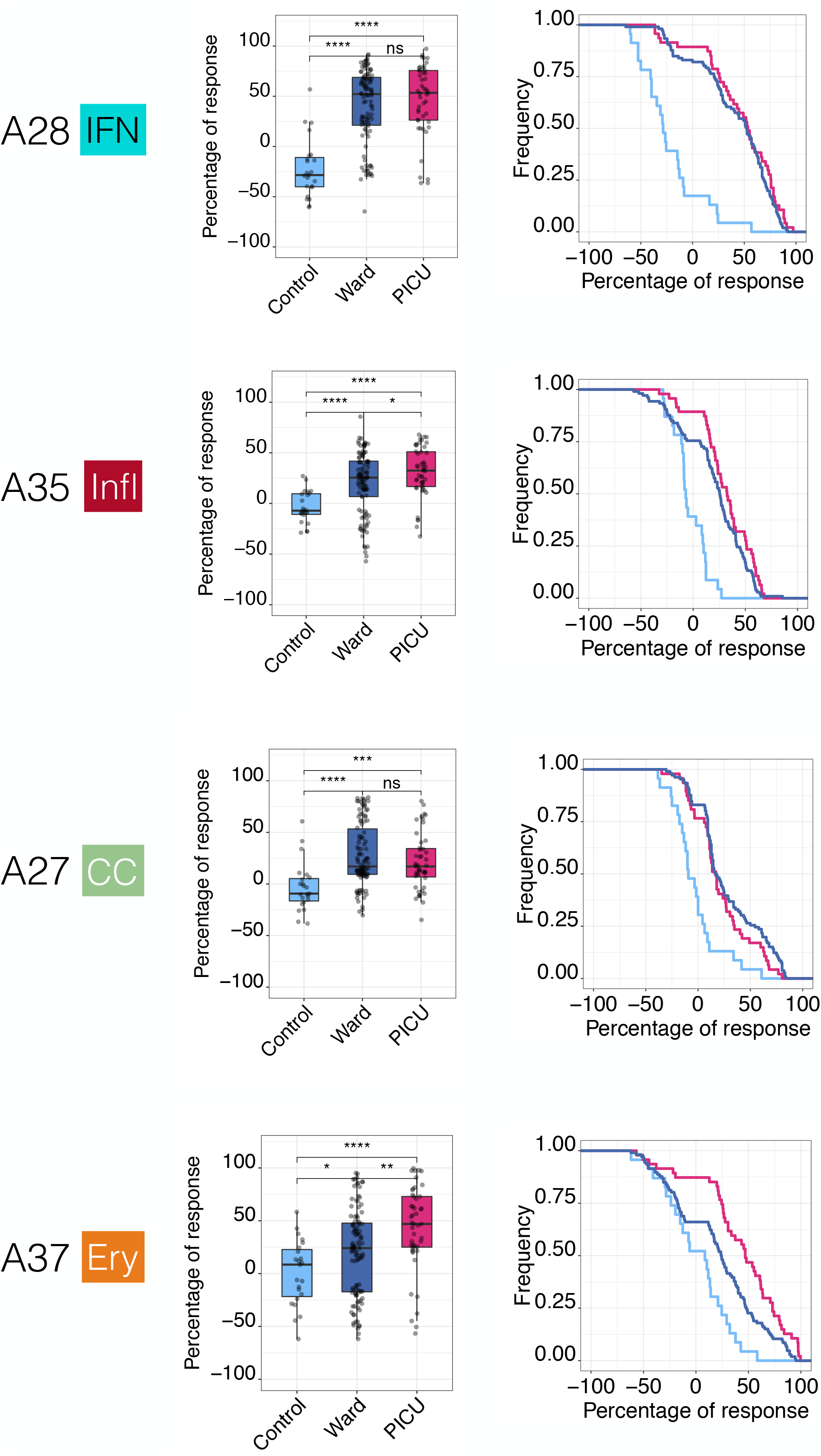
Association of blood transcriptomic traits with RSV disease severity in infants < 4 month of age. Box plots (left) show levels of transcript abundance measured for individual subjects for a given aggregate. This value represents the percentage of transcripts constituting the aggregate that are increased or decreased for an individual compared to the median of the uninfected control group (+100% = all transcripts are increased; −100% = all transcripts are decreased). Individuals are grouped according to health status: uninfected controls, inpatients cared for in the hospital ward or inpatients cared for in the PICU. * p< 0.05, ** p< 0.01, *** p< 0.001. The “Signature survival” curves (right) represent the relative frequency of subjects (y-axis) for whom the percentage response falls at or above a given threshold (x-axis). The percentage response was calculated in the same manner as described for the box plot. Thus, all subjects would have a percentage of response falling between −100% and +100%, as indicated by the curves showing the frequency values of 1 at x = −100%. As the range narrows, the frequency decreases; in most cases a very small proportion of patients have a percentage of response values falling between +90% and +100% (at x = +90%). The separation of the curves is an indication of the differences in the distribution of percentage responses between groups (pale blue = control, dark blue = ward, red = PICU).

Taken together, these findings suggest that our RSV stratification system might be clinically relevant. This conclusion is illustrated by the fact that a high proportion of severe subjects was observed among most phenotypes positive for the Ery trait. This finding might be particularly relevant in infants <4 months–of-age who would otherwise carry a similar risk of developing severe RSV disease when taking age into consideration.

### The RSV “Erythrocyte” signature is shared with melanoma patients and liver transplant recipients

Beyond the question of clinical relevance of these RSV blood transcriptome phenotypes, we next sought to understand their biological significance. For this we relied on several resources. First, we used a web application providing access to a reference collection of module fingerprints for the 16 pathological or physiological states (19) (accessible via: https://drinchai.shinyapps.io/dc_gen3_module_analysis/#; demonstration video: https://youtu.be/y7xKJo5e4).

We generated fingerprint grid plots to compare the changes in transcript abundance in acute influenza and RSV infections (**Figure 8A**). Acute influenza infection highly resembles RSV infection in terms of clinical presentation, especially in infants. As could be expected, the fingerprints of both of these respiratory infections featured a potent interferon signature (modules in aggregate A28; i.e. the IFN trait defined earlier). The modules associated with inflammation (comprised in aggregate A35; Infl trait) were also generally increased in both diseases. However, one of the most marked differences between the influenza and RSV fingerprints concerned the erythrocyte signature (aggregate A37; Ery trait), which was consistently increased in RSV but was unchanged in influenza.

**Figure 8:**
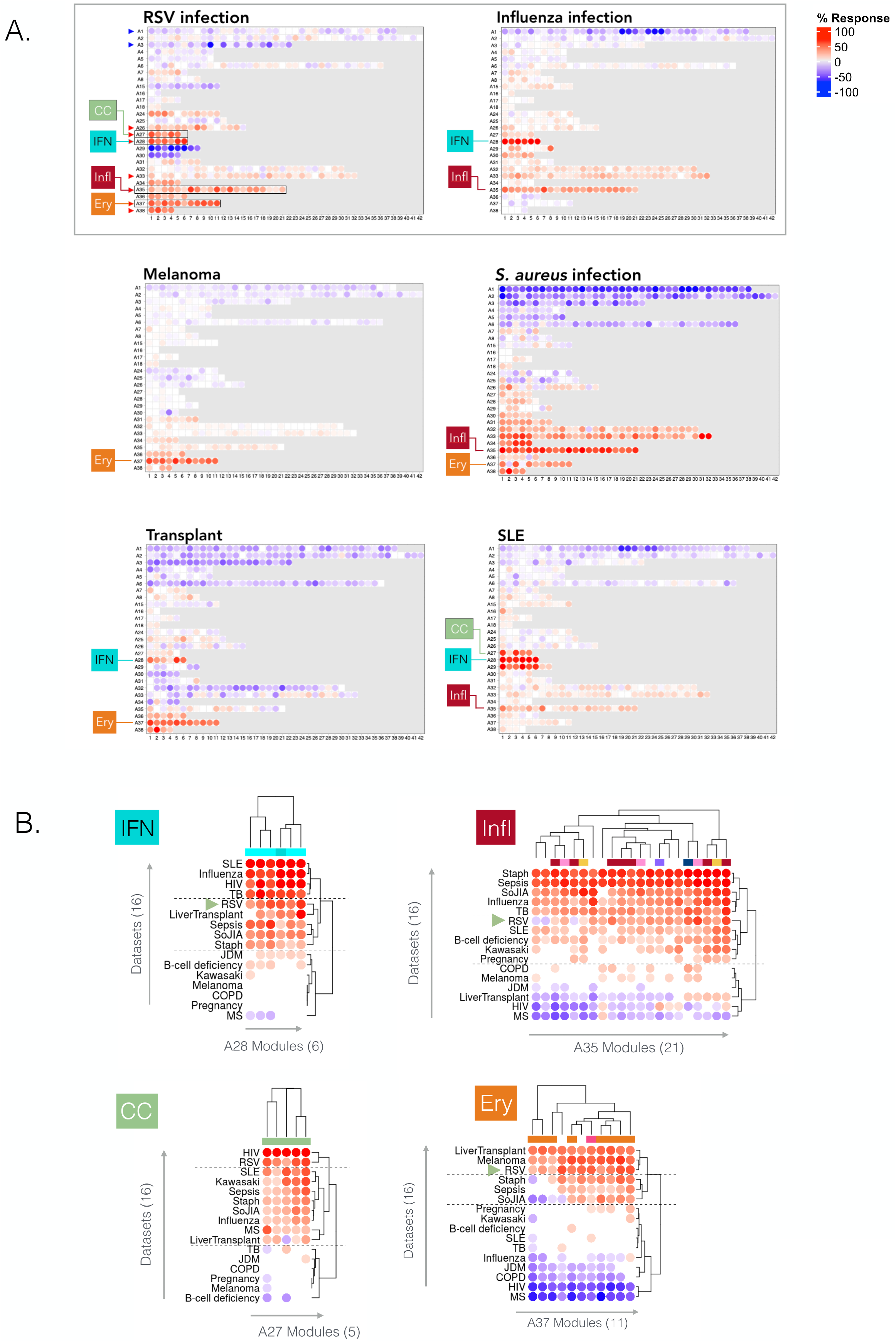
Contextual interpretation of RSV blood transcriptome fingerprints. A. Fingerprint grid plots displaying changes in the levels of transcript abundance in patients with RSV infection, Influenza infection, Systemic Lupus Erythematosus, stage IV melanoma or in liver transplant recipients. The visualization scheme is similar to the one described for the fingerprint grid plot in Figure 1B. The four traits used to molecularly stratify RSV patients are highlighted. Because the position of the modules on the grid is fixed, the color key from Figure 1 can also be used for functional interpretation of the modules from the other rows. The datasets on which the fingerprint maps are based are publicly available under GEO accession ID GSE100150. B. Heatmaps displaying the changes in transcript abundance for modules belonging to four aggregates (columns) across 16 reference datasets. As for the grid plots, an increase and decrease in the abundance of transcripts constituting these modules are shown by a red or blue spot, respectively. The rows (datasets from each disease cohort) and columns (modules) were arranged by hierarchical clustering based on similarities in patterns of transcript abundance. All the plots can be generated and exported via a web application: https://drinchai.shinyapps.io/dc_gen3_module_analysis/# video demonstration https://youtu.be/y7xKJo5e4.

Another fingerprint dominated by an interferon signature was that of systemic lupus erythematosus (SLE), but as was the case for influenza, it did not comprise an “A37/erythrocyte” signature. Among the fingerprints of other reference datasets, those of *Staphylococcus aureus* infection, liver transplantation and metastatic melanoma also showed an elevation in abundance for transcripts constituting modules belonging to the A37 aggregate (**Figure 8A**). The *S. aureus* infection fingerprint showed the highest degree of alteration overall, with widespread changes in transcript levels occurring in most aggregates. This finding contrasts with the fingerprint for metastatic melanoma, which for the most part was quiescent, except for a marked increase in the abundance of genes constituting the A37 modules. The signature observed in liver transplant recipients receiving maintenance chemotherapy was more perturbed than that of melanoma patients, but it was likewise characterized by an increase in the abundance of transcripts constituting the “A37/erythrocyte” modules.

Next, we used the same web application to examine module abundance profiles specifically for the IFN, Infl, CC and Ery traits across all 16 reference datasets (**Figure 8B**). For the IFN trait (A28), RSV clustered among the diseases showing an intermediate level of response, along with liver transplant recipients, patients with systemic onset juvenile idiopathic arthritis (SoJIA), *S. aureus* infection (pediatric) or sepsis caused by various pathogens (adults). Influenza was clustered among diseases showing the highest IFN responses, including other infections such as tuberculosis or HIV, as well as systemic lupus erythematosus (SLE) (**Supplementary Figure 3**). For the Infl trait (A35), RSV clustered again with diseases showing an intermediate response level, and predictably lower than those measured not only in SoJIA, sepsis, and *S.* aureus, but also influenza infection. For the CC trait (A27), the RSV and HIV cohorts formed a cluster with the highest increase in abundance. Finally, for the Ery trait (A37) the RSV cohort was one of only three diseases in the high abundance cluster, along with the melanoma and liver transplant cohorts. We observed increases to a lesser extent in diseases characterized by overt systemic inflammation, such as sepsis or SoJIA, as well as in pregnant women. This trait tended to be decreased in other viral illnesses, such as influenza and HIV infection.

Overall, this contextual interpretation of the dominant traits comprising the RSV signature identified some peculiarities. Notably, the interferon response that, while robust, seemed to be somewhat muted when compared to other viral infections. More strikingly was the atypical overall elevation in the abundance levels associated with the erythrocyte signature. The extent of the observed change was only found in melanoma patients and liver transplant recipients: in these two cohorts, the erythrocyte signature dominated the overall changes observed in the blood transcriptome. Notably, both patient populations relate to states characterized by marked immunosuppression, driven by the disease in the first case and pharmacological treatment aiming at maintaining graft tolerance in the other.

### Expression of transcripts constituting the A37/“erythrocyte” modular signature is restricted to a population of fetal GlyA+ erythroid cells

In our final analyses, we focused our interpretations on the erythrocyte signature (A37). Although we had observed an association with RSV disease severity, we did not ascertain causality. Based on functional profiling results that were run using multiple approaches, we attributed the erythrocyte annotation to 8/11 modules in aggregate A37 (interactive reports available via: https://prezi.com/view/YyQs4WiXSNf0YXE79lfS/). Examining the abundance patterns for the transcripts comprising the A37 modules in reference datasets consisting in isolated leukocyte cell populations provided further insight (**Figure 9;** with additional heatmaps accessible interactively via the weblink provided above). In one such reference dataset contributed by Novershtern et al. (24), the expression of A37 transcripts was narrowly restricted to populations of glycophorin A-positive (GlyA+) fetal erythroid cells. This pattern was irrespective of CD71 marker expression. However, genes comprising A37 modules were not expressed in CD71+ but GlyA-cells. We observed similar expression patterns for modules constituting aggregates A36 and A38, which both comprised one module functionally associated with the erythrocyte annotation (**Supplementary Figure 4**, **Supplementary Figure 5**)

**Figure 9:**
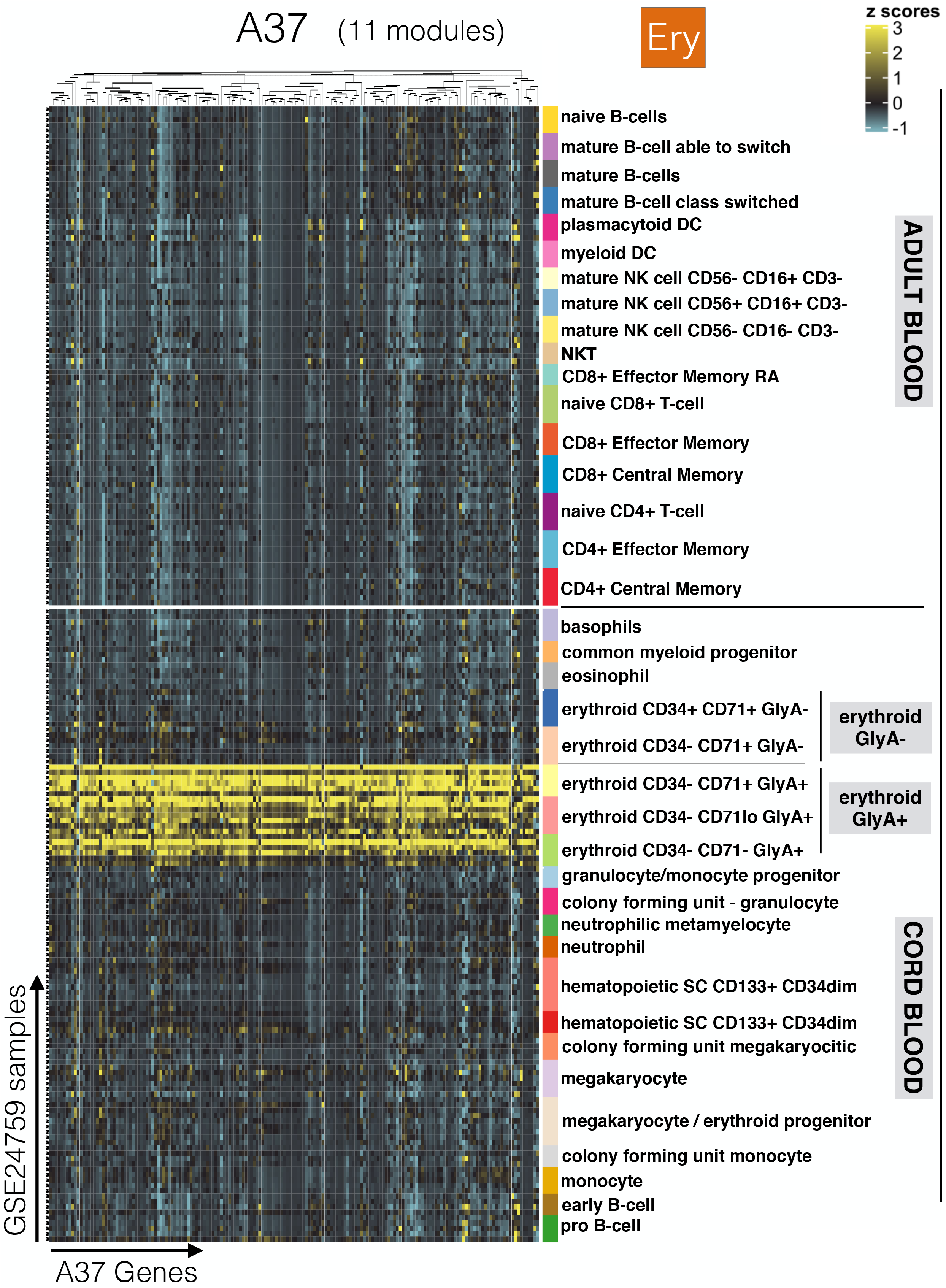
Expression levels of A37 genes across cell populations isolated from human peripheral blood and cord blood. The abundance levels of transcripts comprised in the 11 modules constituting A37 (columns) across blood-cell populations (rows). The dataset is publicly available under GEO accession ID GSE24759 (24). The populations are separated based on whether they were isolated from adult venous blood (top) or from neonate cord blood (bottom). Distinct erythroid cell populations isolated on the basis of cell surface expression of CD34, CD71 and GlyA antigens are also shown.

Erythroid precursors of fetal origin can circulate in the blood of neonates for up to 3-4 weeks following birth. Immunosuppressive properties have been attributed to these populations (25); for example, this cell population confers susceptibility to Listeria infection in neonates (26). However, a possible role for these circulating erythroid cells in the context of RSV infection has not been investigated to date. Others have also described the presence of an erythroid cell population with potent immunosuppressive properties associated with anemia in adults with late stage cancer (27). This finding is consistent with our observation of a prominent A37/erythrocyte signature in the melanoma patients included in our study.

Taken together, these observations support the notion that an increase in A37 transcript abundance is associated with the presence of circulating erythroid cells. These cells might possess immunosuppressive properties and conceivably contribute to the worsening of RSV infection. While these findings might be particularly relevant to young infants, we also observed increases in adult subjects exposed experimentally to the virus (GSE73072: **Figure 1A,Supplementary Figure 1**).

## DISCUSSION

This work has built on earlier studies investigating host responses to RSV via blood transcriptome profiling. The approach we adopted did not focus on identifying sets of classifiers or predictors. Rather, we primarily documented the inter-individual differences among this large consolidated set of patients.

Relying on highly reduced dimensions made it possible to define dominant blood transcriptome phenotypes among this aggregated RSV patient cohort. The four “traits” or signatures that were retained included: interferon (A28 aggregate / IFN trait), inflammation (A35 / Infl), the cell cycle (A27 / CC) and erythrocytes (A37 / Ery). Out of 319 RSV subjects, 199 were distributed in just eight phenotypes out of a possible 81. These dominant phenotypes were all positive for the interferon trait and positive or unchanged for the inflammation and cell cycle traits. The erythrocyte trait status ranged from being increased, unchanged and decreased.

From the standpoint of clinical significance, the phenotypes positive for both the interferon and erythrocyte traits were generally associated with a higher proportion of severe subjects. However, the levels of increase in abundance of interferon-inducible transcripts did not correlate with disease severity. Rather, erythrocyte transcripts showed the strongest association with infection severity. This finding confirmed the association previously described by Mejias et al., which was based on the analysis of one of the datasets comprised in this collection (28). We also observed an association, although to a lesser degree, between the level of increase of transcripts forming an inflammation signature (aggregate A35) and infection severity. Again, this finding is in line with previous data (9,10). Follow-up studies in large patient cohorts are now warranted to validate such classification scheme. Then, the number of traits needed for this classification to be clinically relevant must be determined. Thus far, our analysis suggests that the erythrocyte trait might constitute a valuable risk indicator if testing focuses on a narrowly defined age group (e.g. <4 months-of-age).

Our meta-analysis of available RSV blood transcriptome datasets also yielded several insights relevant to the immune biology of this disease. The interferon signature (A28 / “IFN” trait) is a hallmark of RSV infection and is observed in a wide range of other viral and bacterial infections, as well as autoimmune diseases, as illustrated in the set of 16 blood transcriptome datasets used in our interpretation. Our previous work has suggested that subsets of modules constituting this aggregate are preferentially induced by type I interferons [M8.1, M10.1 (19)]. Consistently, we also observed changes for these modules in patients with RSV infection (**supplementary Figure 3**). Others have also described robust interferon responses as measured via blood transcriptome profiling, in RSV patients (11). The role of type I interferons *per se* is not widely reported, but a recent study describes a dependency on interferon alpha and beta for developing antibody-mediated responses (29). Several reports, however, have identified suppressed interferon gamma responses in the context of RSV infection, especially when compared to the response observed during influenza virus infection (30–33). This finding is also consistent with our observation of a somewhat muted interferon response compared to what we measured in response to not only influenza, but also to HIV or TB infection, as well as in patients with SLE.

We propose that the inflammation signature (A35 / “Infl” trait) is associated with “neutrophil-driven” inflammation, given the preferential expression of its constitutive transcript in neutrophils and induction in patients with sepsis. This information is derived, in part, from a dataset contributed by Linsley et al. that comprised RNAseq profiles of isolated leukocyte fractions (34). The patterns of transcript abundance are available for the modules constituting aggregate A35 via an interactive web presentation (https://prezi.com/view/7Q20FyW6Hrs5NjMaTUyW/). This aggregate was the focus of a recent re-analysis we conducted in which an increase in abundance of A35 transcripts was a dominant feature of the psoriasis blood transcriptome fingerprint (35). We hypothesized in turn that this inflammatory response might be driven more specifically by interleukin-17 (IL17). Indeed, several researchers have found a role for IL17 in the context of RSV infection (36,37) and in one instance specifically indicating the involvement of neutrophils in IL17-mediated antiviral responses (38).

We posit that the cell cycle signature (A27 / “CC” trait) is associated with the expansion of plasmablasts, which are responsible for antibody production. Indeed, modules in this aggregate comprised an overabundance of genes involved in the cell cycle, such as cyclins. One of the modules also comprises several genes expressed by plasmablasts (M12.15: CD38, IGJ, TNFRSF17) (39–41) that are markedly expressed between 7 and 14 days after the administration of trivalent influenza or pneumococcal vaccines (42). In the context of this study, changes in abundance of these markers was confirmed by flow cytometry to be correlated with the presence of these antibody producing cells. These levels also correlated with antibody titers measured four weeks post-vaccination. These cell populations are also expanded during the course of RSV infection. Habibi et al. reported a peak 10 days after experimental exposure to RSV and an correlation with the levels of neutralizing antibody developed by the individuals (43).

Consistent with our earlier findings (28), the erythrocyte signature (A37 / “Ery” trait) was most strongly associated with severity. However, such an association has not been reported in other studies comprising the consolidated dataset collection. We putatively link this signature with the presence of circulating erythroid cell precursors, based on the restriction of A37 transcripts in a reference transcriptome dataset to fetal erythroid cells (**Figure 9**). Erythroid precursors would normally be found in the bone marrow, but cells originating from the fetal liver do persist in infants in the circulation for a few weeks after birth (44). In adults, extramedullary erythropoiesis is observed in the spleen and liver, and occurs under various circumstances, including anemia, pregnancy, severe infection or chronic stress (45,46). We hypothesize that circulating erythroid cells (CECs) associated with this signature might have immunosuppressive functions during an RSV infection. Thus, CEC-mediated immunosuppression would in turn drive a worsening of disease and severity in patients with RSV infection. This assertion is supported by various lines of evidence. First, Elahi et al. have described a wide range of mechanisms conferring immunosuppressive properties to this cell population, including via soluble factors (such as arginase, TGFbeta, reactive oxygen species) or cell surface receptors (such as PD1/PDL-1 and VISTA) (25). Second, we also observed a marked increase in A37 transcript abundance in metastatic melanoma and liver transplant recipients under maintenance therapy. Both of these states are characterized by marked immunosuppression and were categorized in the high abundance profile cluster for A37. Of the 14 other cohorts in this reference collection only RSV was present in this same cluster. Others have recently described immunosuppression exerted by CECs in patients with late stage cancer: these cells were found to be at least in part responsible for the impaired T-cell responses observed in this patient population (27). Third, the RSV literature provides indications of this virus’ ability to subvert the immune response (4). While the possibility of an involvement of CECs in this immune modulation of the response to RSV is novel, the contribution of the hyporesponsiveness of the neonatal immune system as an underlying factor to progression to severe RSV infection has been clearly outlined (47). A key question could thus center on the possible contribution of CECs to the reduced competence of the neonatal immune system. Indeed, results obtained by Elahi et al. in animal models indicate that CEC depletion can restore neonatal immune responsiveness and confer resistance to Listeria infection (26).

The synthesis that we conducted here builds upon and extends earlier findings. It also identifies new avenues of investigations. Notably, it points to the potential relevance of blood transcriptional phenotyping for stratification of patients with RSV infection. More specifically, a signature putatively attributed to immunosuppressive erythroid cells was found to be associated with clinical severity, even in homogenously younger patients. Clinical relevance of such candidate biomarker signature would need to be assessed next. Further investigation of circulating erythroid cells population in the context of RSV infection are also warranted by these findings. One of the central questions being whether these cells merely accompany clinical worsening of the disease or constitute one of its drivers. More generally, this work also highlights the need for follow on large-scale blood transcriptome profiling studies of responses to RSV patients, especially over multiple time points. Coordination and cooperation between the groups that may engage in such endeavors would also prove beneficial for generating large inter-operable blood transcriptome dataset collections.

## ACKNOWLEDGEMENTS

This work is based in part on analyses conducted during a training workshop supported by Inflammation-Immunopathology-Biotherapy Department (i2B), AP-HP, Pitié-Salpêtrière Hospital, LabEx Transimmunom (ANR-11-IDEX-0004-02) and RHU iMAP grants to DK and organized with the help of Caroline Aheng and Sophie Miller. Sidra Medicine is a member of the Qatar Foundation for Education, Science and Community Development. We would like to acknowledge Insight Editing London for editing the manuscript prior to submission.

## CONTRIBUTIONS

Conceptualization: DR, DC, DB, EM, DK, MA. Data curation and validation: DR, MT, MG, BK, OK, SA, FM. Visualization: DR, DC, OK, SA, FM. Analysis and interpretation: DR, OK, SH, FM, AM, OR, EM, DK, DC. Writing of the first draft: DR, DC. Funding acquisition: DK, DC. Methodology development: DR, DC. Writing – review & editing: DR, MA, MT, MG, BK, OK, SH, FM, AM, OR, DB, EM, DK, DC. The contributor’s roles listed above follow the Contributor Roles Taxonomy (CRediT) managed by The Consortia Advancing Standards in Research Administration Information (CASRAI) (https://casrai.org/credit/).

---

**Supplementary Figure 1:**
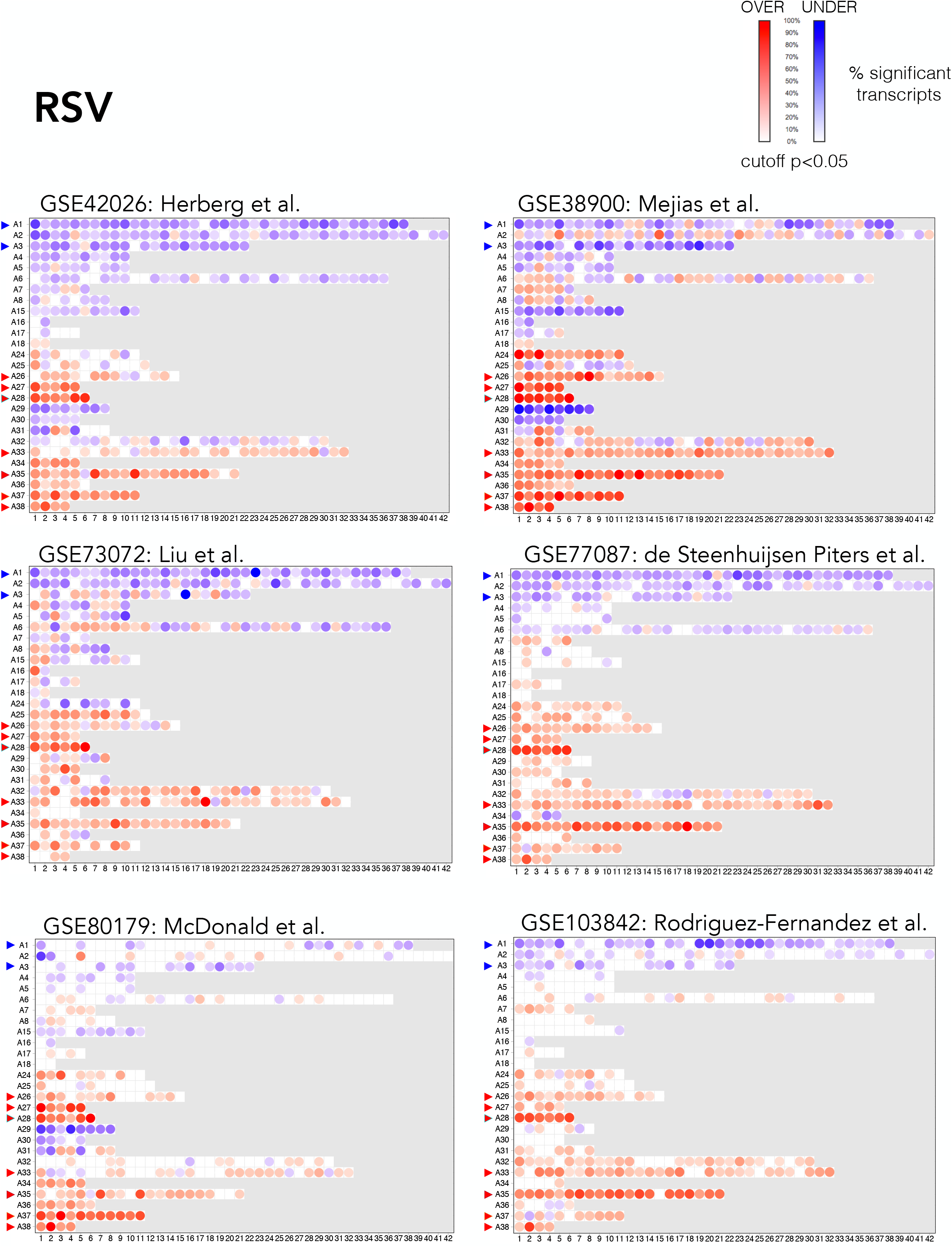
Fingerprint grid maps of modular repertoire changes in six independent RSV blood transcriptome datasets. The visualization scheme is similar to the one described for the fingerprint grid map in Figure 1B. Because the position of the modules on the grid is fixed, the color key from Figure 1 can also be used for functional interpretation. The maps were generated from multiple independent RSV blood transcriptome datasets that are available in the NCBI Gene Expression Omnibus (5,6,20–22,28).

**Supplementary Figure 2:**
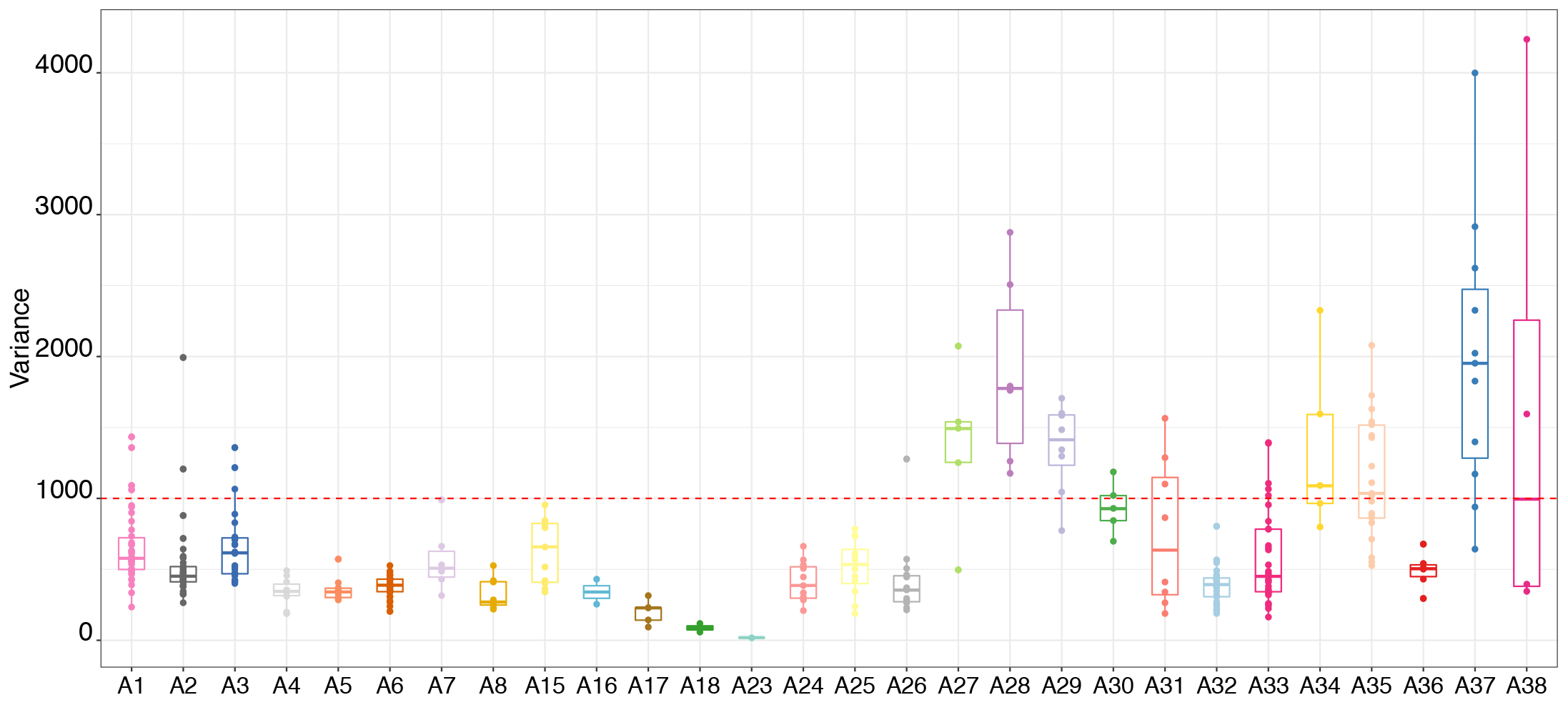
Inter-individual variance of summarized module-level abundance values. The variance in the percentage of the transcript response across 319 RSV patients was calculated across 382 modules. The box plots display those values with each module being grouped in its respective module aggregate, from A1 to A38.

**Supplementary Figure 3:**
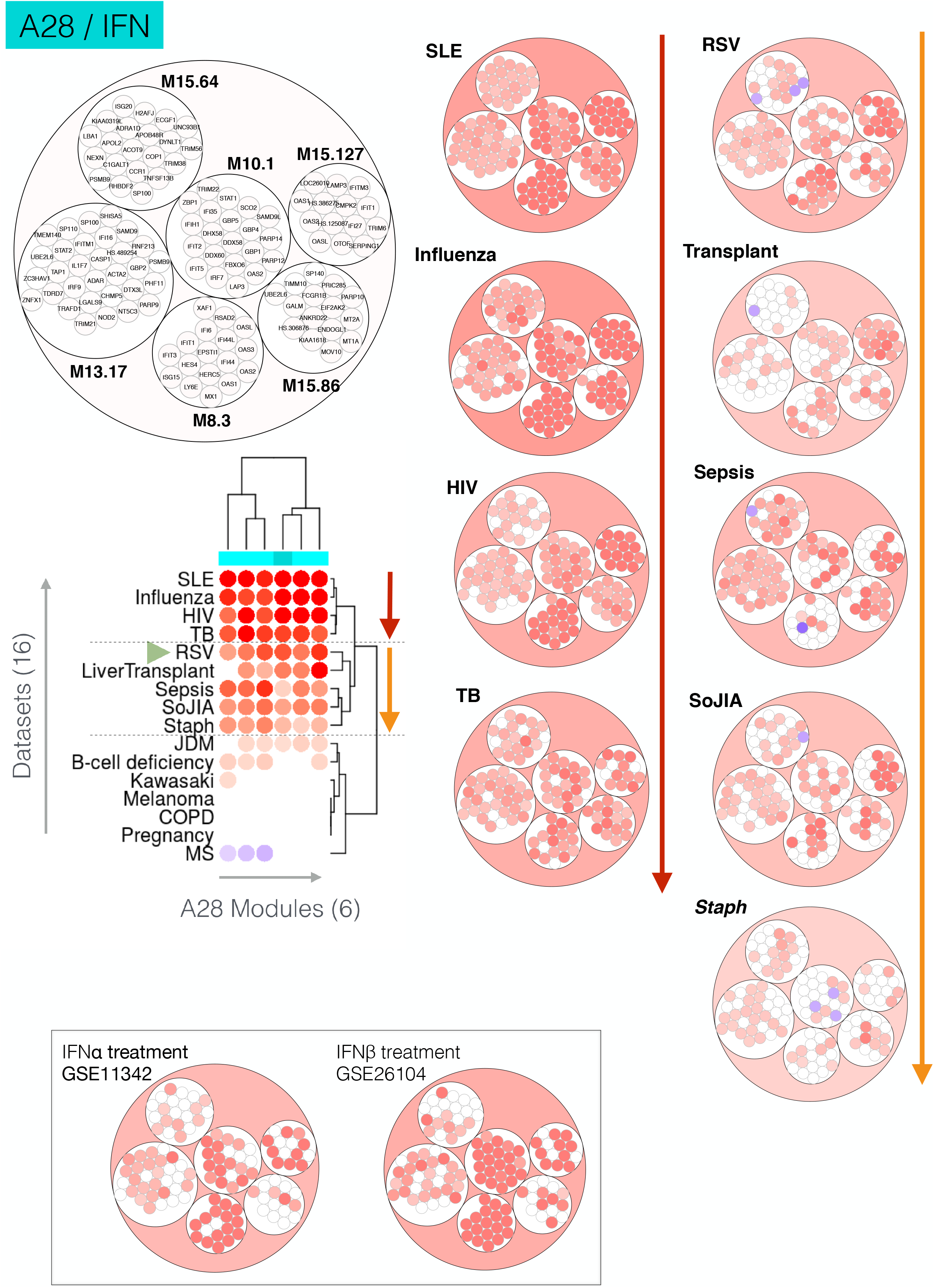
Patterns of abundance of six A28 modules across several disease and physiological states. Each column on the heatmap corresponds to one of six modules constituting the A28 aggregate. Each row corresponds to one of 16 reference datasets. Red spots on the heatmap indicate an increase in abundance of the transcripts constituting a given module for a given dataset. A blue spot indicates a decrease in abundance. No color indicates no change. Disease or physiological states were arranged based on similarity in patterns of aggregate activity. The circle (top left) is a representation of the six modules constituting aggregate 28, and of the transcripts constituting each of the modules. Some genes on the Illumina BeadArrays can map to multiple probes, which explains the few instances where the same gene can be found in different modules. The smaller circles (right) represent changes in transcript abundance of individual transcripts for two of the disease clusters. The first cluster (red arrow) comprises four diseases with the highest degree of increase in transcript abundance. The second cluster (orange arrow) comprises five diseases with an intermediate level of increase in abundance. The insert (bottom) shows two circles representing changes in abundance in patients with hepatitis C treated with IFNα(51), and patients with multiple sclerosis treated with IFNβ(52). The corresponding NCBI GEO accession IDs are indicated for each.

**Supplementary Figure 4:**
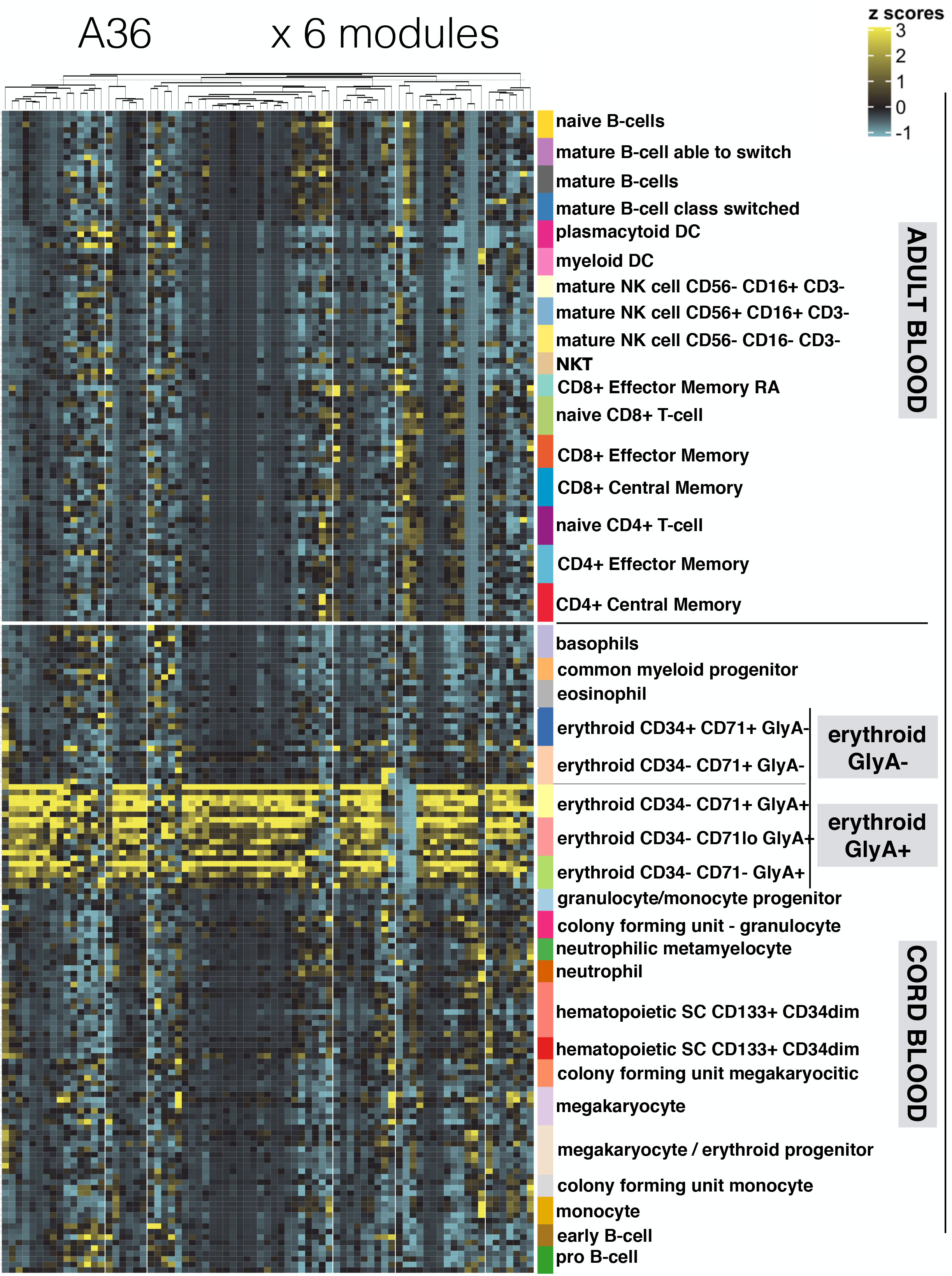
The expression levels of A36 genes across the cell populations isolated from human peripheral blood and cord blood. The abundance levels of transcripts comprised in the 11 modules constituting aggregate A36 (columns) across blood cell populations (rows). The dataset is publicly available under GEO accession ID GSE24759 (24). The populations are separated based on whether they were isolated from adult venous blood (top) or from neonate cord blood (bottom). Distinct erythroid cell populations isolated on the basis of cell surface expression of CD34, CD71 and GlyA antigens are shown.

**Supplementary Figure 5:**
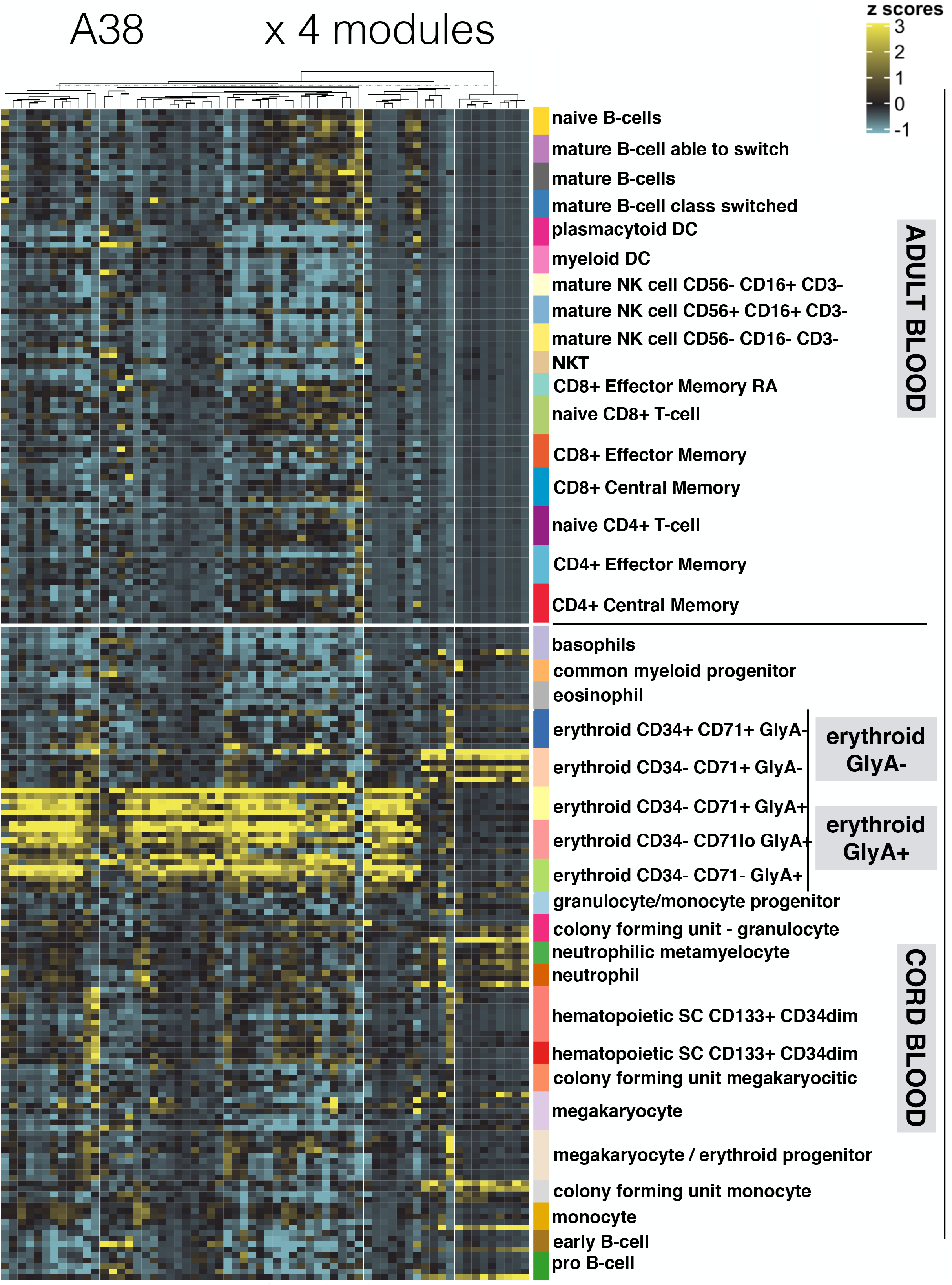
The expression levels of A38 genes across the cell populations isolated from human peripheral blood and cord blood. The abundance levels of transcripts comprised in the 11 modules constituting aggregate A38 (columns) across blood cell populations (rows). The dataset is publicly available under GEO accession ID GSE24759 (24). The populations are separated based on whether they were isolated from adult venous blood (top) or from neonate cord blood (bottom). Distinct erythroid cell populations isolated on the basis of cell surface expression of CD34, CD71 and GlyA antigens are shown.

## METHODS

### Selection of public blood transcriptome datasets

Datasets deposited in the NCBI Gene Expression Omnibus, GEO, were used in this meta-analysis. Accession IDs along with descriptive information and references can be found in **Table 1**. A reference dataset, which consisted of transcriptome profiles derived from adult blood cell populations and cord blood was also used to support the functional interpretation of our findings. This dataset was contributed to the GEO by Novershtern et al. with accession ID GSE24759 (24).

### Module repertoire construction

The construction of a transcriptional module repertoire for blood transcriptome analyses has been described previously (48,49). The version that was used in this study is the third one developed by our group and is the object of a separate publication (available on a pre-print server (19)). Briefly, the approach consists of identifying sets of co-expressed transcripts for a given biological system (in this case blood) and across a wide range of disease or physiological states (perturbations of steady state). In this case, co-expression was determined based on patterns of co-clustering observed for all gene pairs across a collection of 16 reference datasets. These datasets encompassed viral and bacterial infectious diseases (HIV, influenza, RSV, Melioidosis, *Staphylococcus aureus*, Tuberculosis) as well as several inflammatory or autoimmune diseases (systemic lupus erythematosus, multiple sclerosis, chronic obstructive pulmonary disease, Kawasaki disease, juvenile dermatomyosistis, systemic onset juvenile idiopathic arthritis), B-cell deficiency, liver transplantation, stage IV melanoma and pregnancy. The overall collection comprised 985 blood transcriptome profiles. A weighted co-expression network was built on the basis of co-clustering patterns that were obtained. Here, the weight of the nodes connecting a gene pair being based on the number of times co-clustering was observed, thus ranging from a weight of 1 (where co-clustering occurs in one of 16 datasets) to a weight of 16 (where co-clustering occurs in all 16 datasets). Next, this network was mined using a graph theory algorithm (identification of cliques and paracliques) to define a subset of densely connected gene sets that constituted our module repertoire. Overall, 382 modules were identified via this process, encompassing 14,168 transcripts. A supplemental file including the definition of this module repertoire along with the functional annotations is available from a companion publication.

### Constitution of module aggregates

To maintain the number of variables within a manageable number and to facilitate data interpretation, a second tier of clustering was performed to group the modules into “aggregates”. This was achieved by segregating the set of 382 modules according to the patterns of transcript abundance across the 16 reference datasets that were used for module construction. This segregation resulted in the formation of 38 aggregates, each comprising between one and 42 modules. The second level of granularity that was thus obtained was used to define distinct RSV blood transcriptome phenotypes and as a basis for the fingerprint grid plot representation (see **Figure 1** and **Figure 7**). With such grids, the first vertical reading of the fingerprint grid provides an overview of the changes in transcript abundance observed among module aggregates, while the horizontal reading provides the changes observed within an aggregate and across modules.

### Module repertoire analysis workflow

The modular analysis was performed using 14,168 transcripts. The fold change was computed using gene expression data prior to log2 transformation. For group comparisons, a paired t-test was performed on the log2-transformed data [Fold change (FC) cut off = 1.5; FDR cut off = 0.1]. For individual-patients analyses, each sample was compared to the mean value of control samples in each dataset. The cut off comprised an absolute FC >1.5 and a difference in gene expression level >10. The results for each module analysis are reported as the percentage of its constitutive transcripts for which the abundance was increased or decreased. Because gene sets are selected based on the co-expression observed in blood, the changes in abundance within a given module tend to be coordinated and the dominant trend is therefore selected (the greater value of the percentage increased vs percentage decreased). A module was considered to be “responsive” when the proportion of differentially expressed transcripts (as defined above) was >15%.

### Data visualization

The results were visualized in a fingerprint format, either as a grid plot (group level, **Figure 1**) or as a heatmap (individual level, **Figure 5**), using the same illustrative RSV dataset. For each module, the percentage of increased transcripts is represented by a red spot and the percentage of decreased transcripts is represented by a blue spot. The largest of the two values was retained for visualization. In the grid format (**Figure 1**), the position of each module is fixed. A row of modules corresponds to a “module aggregate”, which as described above, is a set of modules following a similar pattern of activity across the 16 input datasets corresponding to different disease or physiological states. A few “aggregates” comprised only a single module and thus are not shown on the grid. The fingerprint grid plots were generated using “ComplexHeatmap” (50).

For the heatmaps (**Figure 2, Figure 5, Figure 7B)**, each row corresponds to a module and each column to a sample. The columns and rows are arranged based on similarities in the patterns of module activity. Filters can be applied to remove modules that show only low levels of activity across the samples or to retain only the modules associated with functional annotations.

