## Supplementary figures and images for "Identification of erythroid cell positive blood transcriptome phenotypes associated with severe respiratory syncytial virus infection"

### Supplemental File 1

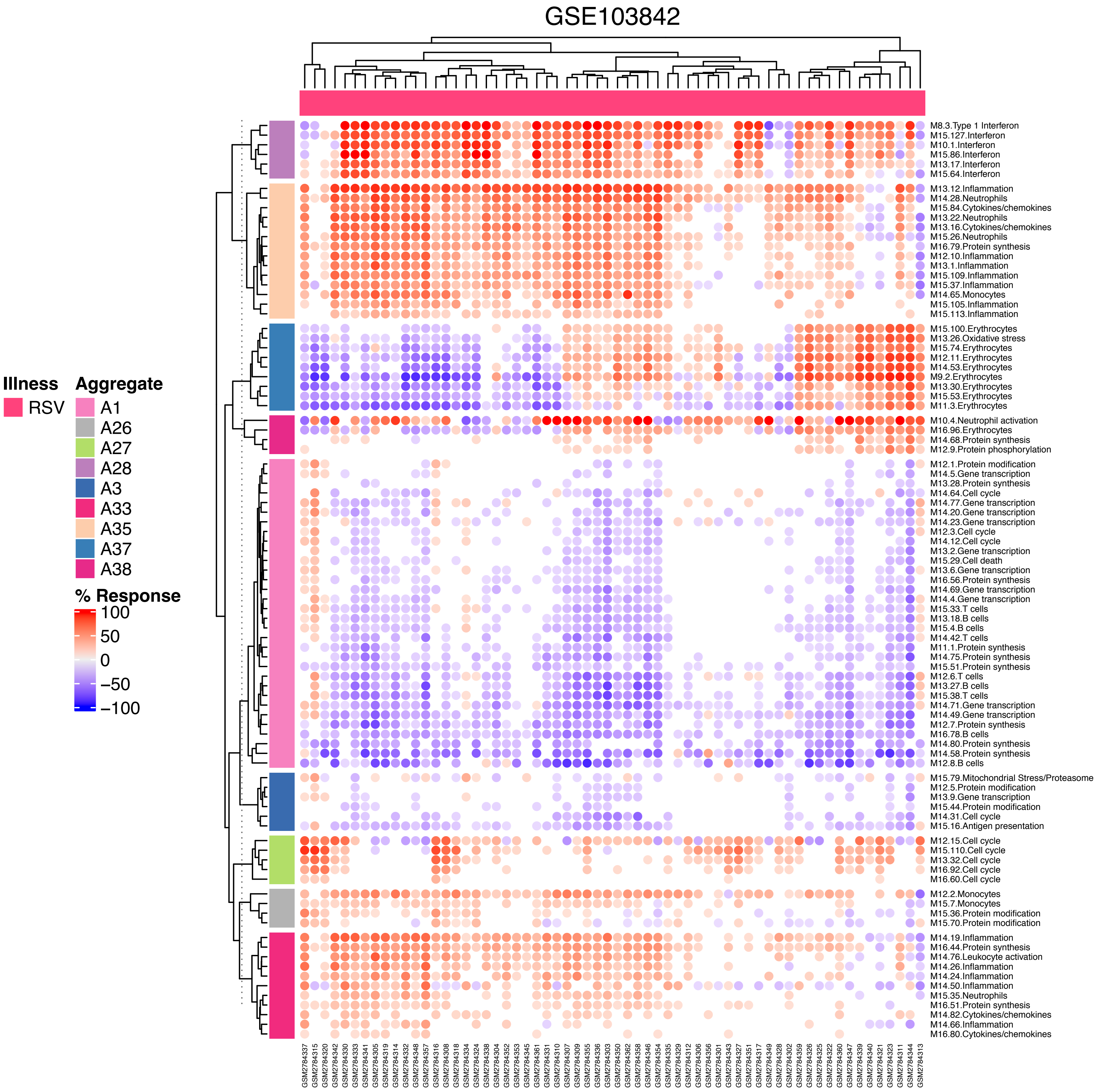

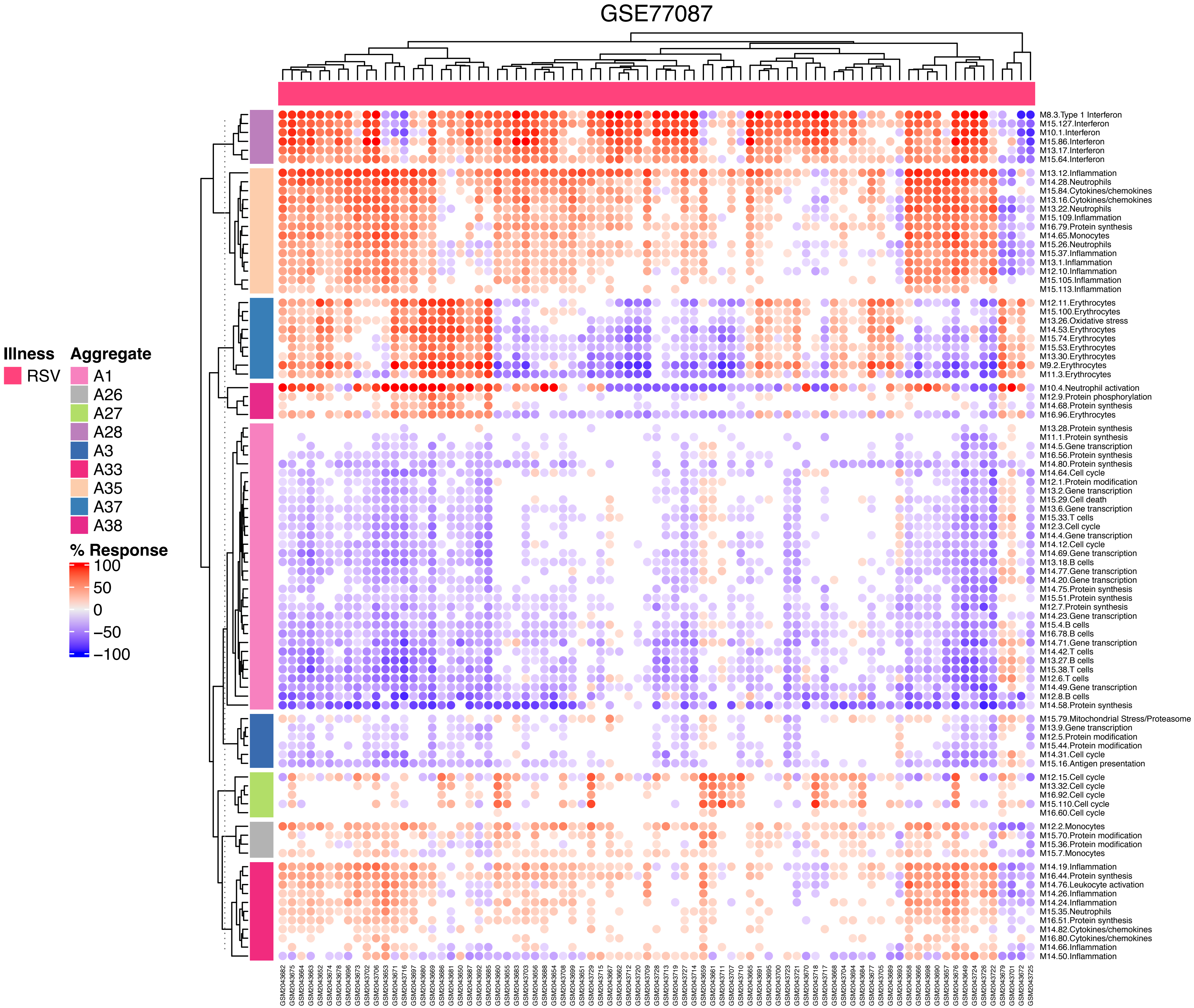

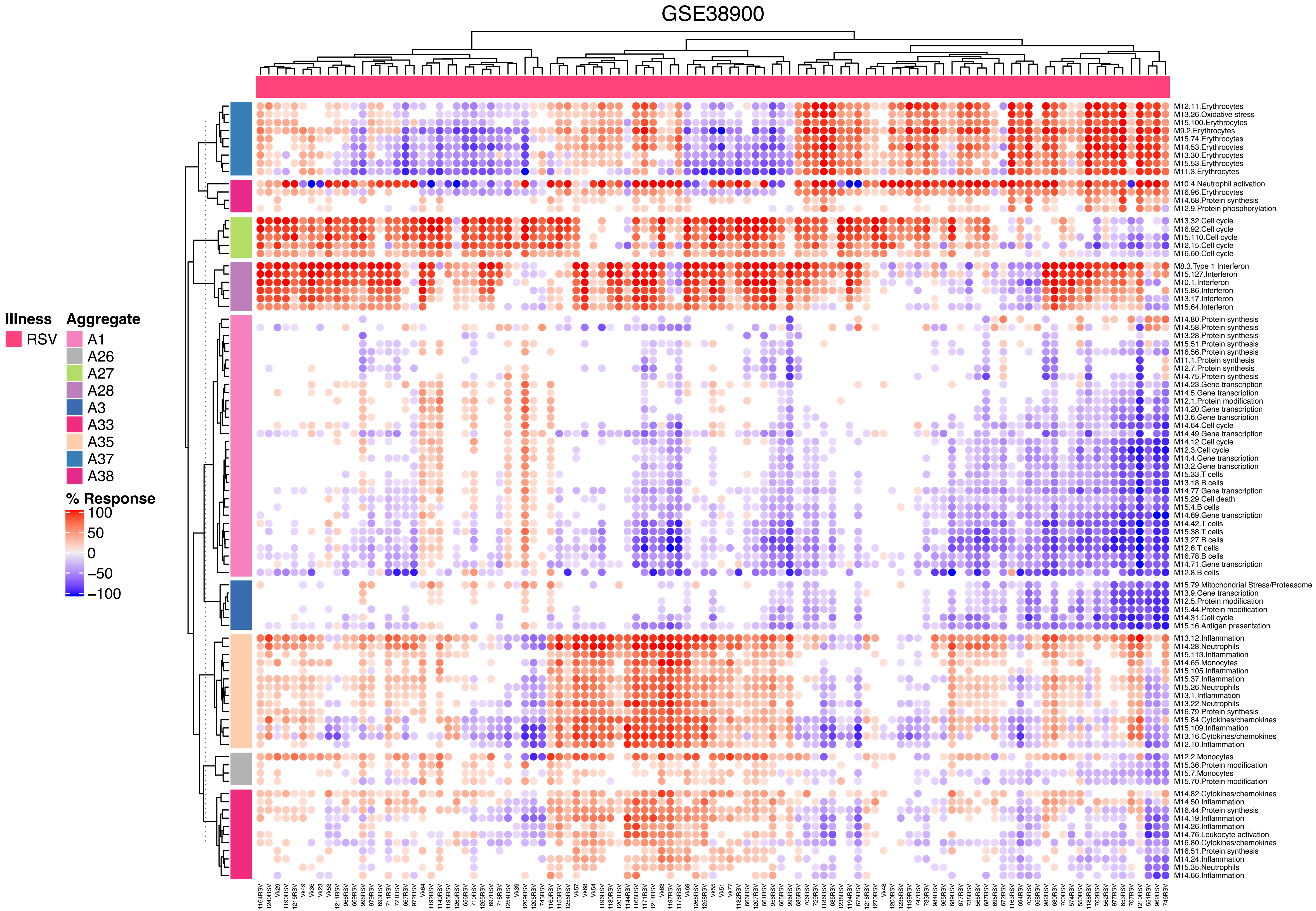

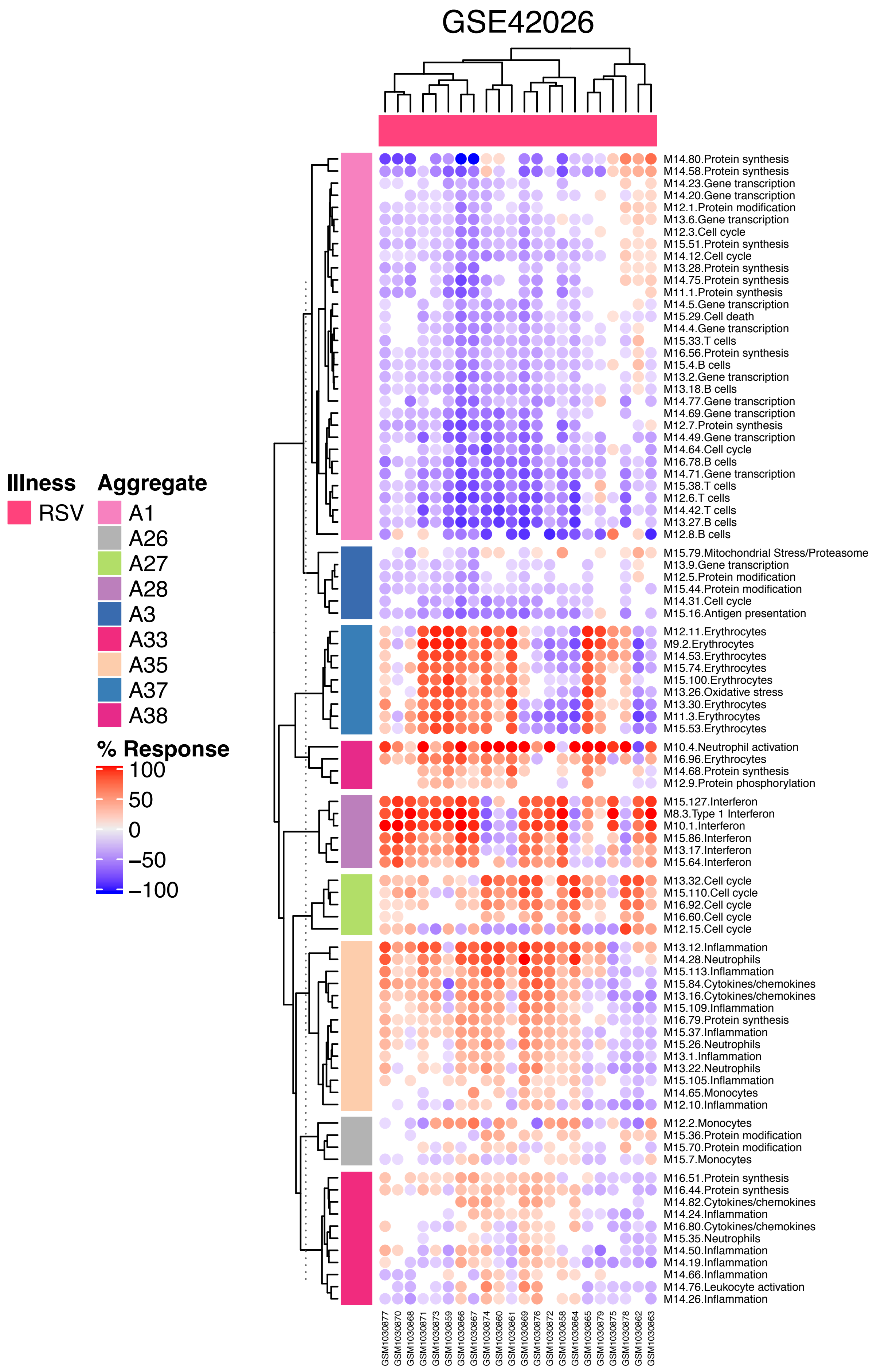

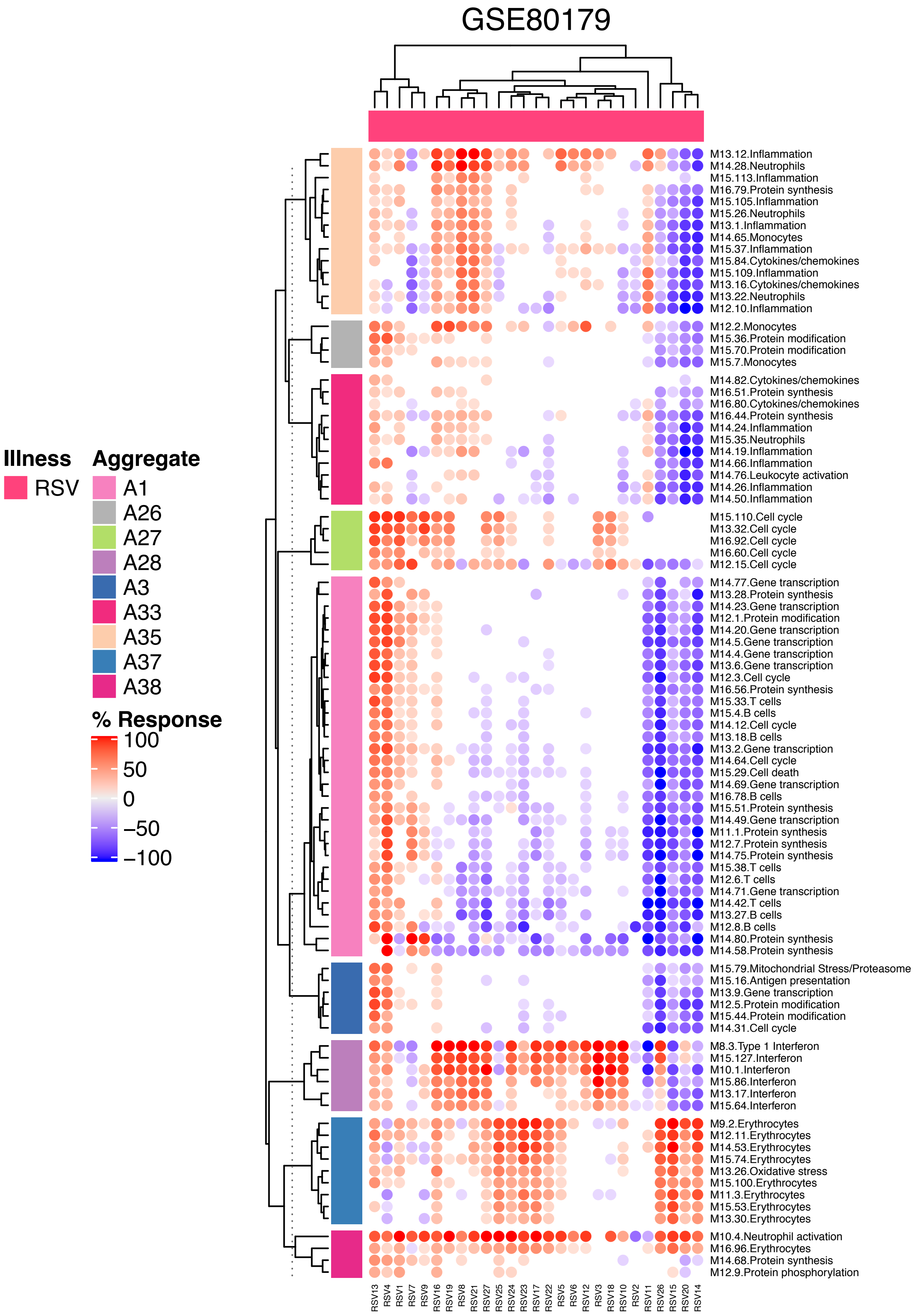

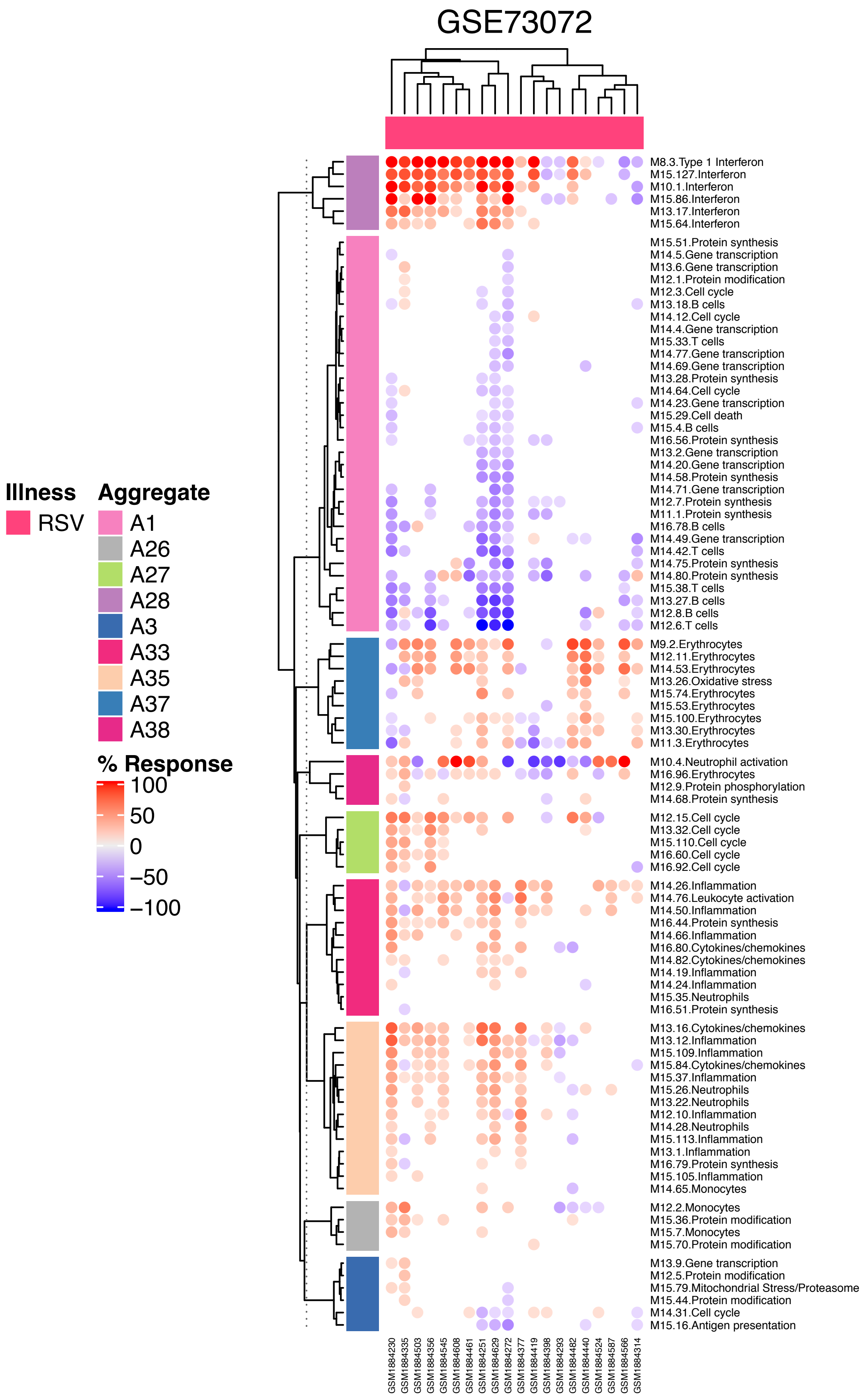
